# *TCF-7* is dispensable for anti-tumor response but required for persistent function of mature CD8 T cells

**DOI:** 10.1101/2021.03.22.436494

**Authors:** Rebecca Harris, Mahinbanu Mammadli, Shannon Hiner, Liye Suo, Qi Yang, Jyoti Misra Sen, Mobin Karimi

## Abstract

T Cell Factor-1, encoded by *TCF-7,* is a transcription factor that plays an essential role during T cell development and differentiation. In this manuscript we utilized a pre-clinical model provided evidence that *TCF-7* is dispensable for the anti-tumor response, and that *TCF-7* suppresses key transcriptional factors Eomes and T-bet and molecules responsible for peripheral CD8 T cell cytolytic function. We discovered that *TCF-7* regulates NKG2D expression on naïve and activated mouse CD8 T cells, and that peripheral CD8 T cells from *TCF-7* cKO utilize NKG2D to clear tumor cells.

We also provide evidence that *TCF-7* regulates key signaling molecules, including LCK, LAT, ITK, PLC-γ1, P65, ERKI/II, and JAK/STATs required for peripheral CD8 T cell persistent function. Our data transcriptomic and protein data uncovered the mechanism of how *TCF-7* impacting peripheral CD8 T cell inflammatory cytokine production, CD8 T cell activation, and apoptosis. Our pre-clinical model showed that CD8 T cells from *TCF-7* cKO mice did not cause GVHD, but effectively cleared primary tumor cells.

## Introduction

T Cell Factor-1 (*TCF-7*), a major T cell developmental transcription factor, is involved in the Wnt signaling pathway, and is critical for T cell development as well as activation (Escobar et al., 2020; Ma et al., 2012). Dysfunction of the Wnt/β-catenin/*TCF-7* signaling pathway leads to immune deficiency or autoimmunity (Shi et al., 2016). It is well known that *TCF-7* is involved in the regulation of cell proliferation and survival during later T cell development (Kim et al., 2020). In the absence of *TCF-7*, T cell development is completely blocked after early thymocyte progenitor (ETP) cells, which suggests that *TCF-7* is also important in T cell lineage specification (Germar et al., 2011; Weber et al., 2011). Several studies have shown that *TCF-7* is critical for controlling viral infection (He et al., 2016; Im et al., 2016; Utzschneider et al., 2016). Furthermore, *TCF-7* + CD8+ T cell have self-renewal capacity, while CD8+ T cells lacking *TCF-7* do not (Utzschneider *et al*., 2016; Wu et al., 2016). Overwhelming evidence has suggested that *TCF-7* is critical for CD8 T cell persistence and capacity to control viral infection (Kurtulus et al., 2019; Miller et al., 2019; Siddiqui et al., 2019; Wu *et al*., 2016). While investigating the role of *TCF-7* in viral infection, single cell RNA sequencing data uncovered that CD8 T cells expressing higher levels of *TCF-7* are quiescent and tissue resident, while CD8 T cells expressing higher levels of *TCF-7* in periphery are highly proliferative, and that *TCF-7* negative T-bet^hi^ CD8 T cells are transitory effector cells. *TCF-7*^−^ Eomes^hi^ CD8 T cells are not proliferative and express higher levels of immune checkpoint receptors, but maintain effector function (Beltra et al., 2020; LaFleur et al., 2019; Zander et al., 2019). Studies have also reported that *TCF-7*+ CD8 T cells with stem-like abilities expresses low levels of PD-1 and Tim checkpoint receptors, further providing evidence that *TCF-7* is required for CD8 T cell formation and persistent function (Kurtulus *et al*., 2019; Siddiqui *et al*., 2019; Utzschneider *et al*., 2016). Studies have demonstrated that dysfunctional virus-specific CD8 T cells transplanted into naïve mice give rise to *TCF-7*-negative CD8 T cells (Utzschneider et al., 2013). However, tumor-specific CD8 T cells transferred to naïve mice might give rise to *TCF-7*+ CD8 T cells, demonstrating the differential role of *TCF-7* in viral infection and tumor (Schietinger et al., 2016). These differences could be due to the differences between microenvironments in viral infection and in tumors (Philip et al., 2017). Whether *TCF-7*+ cells will give rise to *TCF-7*+ or *TCF-7*- cells will be dependent on internal and external signaling. Some studies have shown that both the long (p45) and the short (p33) *TCF-7* isoform are expressed by CD8 T cells that will give rise to stem-like CD8 T cells during viral infection, and it has been shown that the long isoform of *TCF-7* is capable of promoting stem-like CD8 T cell formation during viral infection by regulating genes like CD127, CXCR5 and cMyb (Chen et al., 2019). Currently, the role of long and short *TCF-7* isoforms is largely unknown.

Studies of T cells as immunotherapy in both human and mice showed that for a superior anti-tumor response, less differentiated cells are more favorable than a more differentiated subset of CD8 T cells (Gattinoni et al., 2011; Im *et al*., 2016; Lugli et al., 2013). Ideal CD8 T cells for immunotherapy have been shown to exhibit stem-like abilities (such as those obtained by inducing *ex vivo* cell growth with IL-17, IL-15, IL-21) and higher expression of *TCF-7*, Eomes, and Bcl6 (Cieri et al., 2013; Cui et al., 2011). Thus, the suitability of *TCF-7* as a target for immunotherapy to clear viral infection and cancer might also indicate considerable consequences for autoimmune diseases.

To study the role of *TCF-7* in a clinically relevant model, we utilized allogeneic hematopoietic stem cell transplantation (allo-HSCT). In allo-HSCT, mature peripheral donor T cells found in the graft become alloactivated upon recognition of host HLA as non-self. These T cell activities help to clear residual malignant cells, which is called the graft-versus-leukemia effect (GVL) (Balassa et al., 2019; Giralt and Bishop, 2009; Hall and Shenoy, 2019). On the other hand, alloactivated T cells also target healthy recipient tissues, an effect known as graft-versus-host disease (GVHD) (Mavers and Bertaina, 2018). We used a unique mouse strain which has deletion of *TCF-7* in mature T cells, rather than a global deletion (Xing et al., 2016). This *TCF-7* flox/flox x CD4cre mouse experiences deletion of *TCF-7* in all T cells at the double-positive phase of development, when all T cells express CD4 (Wang et al., 2019b). This allows us to overcome the severe T cell developmental defect that occurs in global *TCF-7* deletion, as *TCF-7* is critical for the double-negative stage of development (Yang et al., 2019).

Using a mouse model of GVHD and GVL following allo-HSCT, we were able to study all of the major T cell functions, as well as phenotype, clinical outcomes, and gene expression, in a single model. In this model, we transplanted CD8 T cells from either WT or *TCF-7* cKO mice into irradiated BALB/c mice (H2K^b^→H2K^d^) (Mammadli et al., 2021a; Mammadli et al., 2021c; Mammadli et al., 2021d). Using allogenic pre-clinical model, we have shown that CD8 T cells from *TCF-7* cKO effectively clear tumor cells without inducing GVHD by producing significantly less inflammatory cytokines as proinflammatory cytokines are con sidered the hallmark of allo-immunity (Ju et al., 2005; Seif et al., 2017). Our data also uncovered that CD8 T cells from *TCF-7* cKO mice cause significantly less tissue damage in GVHD target organs (Bleakley et al., 2012; Breems and Lowenberg, 2005). Molecular analysis showed that CD8 T cells from *TCF-7* cKO mice significantly showed reduction in key molecules required for CD8 T cell persistent function (Germar *et al*., 2011; Giralt and Bishop, 2009; Gounari and Khazaie, 2022). CD8 T cells from *TCF-7* cKO mice exhibited innate memory-like phenotype by upregulating CD122, CD44, and effector and central memory phenotypes in mature CD8 T cells critical for GVHD development (Dutt et al., 2011; Huang et al., 2013). We also uncovered that CD8 T cells from *TCF-7* cKO mice significantly upregulated Eomes and T-bet, two downstream transcription factors which are known to be involved in GVL (Mammadli *et al*., 2021a; Weeks et al., 2021; Zhou et al., 2010). Our data demonstrated that naïve CD8 T cells from *TCF-7* cKO mice upregulated NK cell type 2 receptor (NKG2D). NKG2D, encoded by *Klrk1*, is an activating cell surface receptor that is predominantly expressed on Natural killer cells (Larson et al., 2006; Wensveen et al., 2018). While naïve human CD8^+^ T cells express NKG2D, in mice CD8 T cells only upregulate NKG2D upon activation (Hu et al., 2016; Maasho et al., 2005). Upregulation of Granzyme B on CD8 T cells from *TCF-7* cKO mice was also observed. The loss of *TCF-7* also led to upregulation of NK cell type 2 receptor (NKG2D). NKG2D, encoded by *Klrk1*, is an activating cell surface receptor that is predominantly expressed on Natural killer cells (Larson *et al*., 2006; Wensveen *et al*., 2018). NKG2D is abundantly present on all NK cells, NKT cells, and subsets of γδ T cells (Stojanovic et al., 2018). While naïve human CD8^+^ T cells express NKG2D, in mice CD8 T cells only upregulate NKG2D upon activation (Hu *et al*., 2016; Maasho *et al*., 2005). Upregulation of Granzyme B on CD8 T cells from *TCF-7* cKO mice was also observed. Our molecular and animal data were confirmed by transcriptomic analysis.

Altogether, our data demonstrate that loss of *TCF-7* in mature murine CD8 T cells enhanced Eomes and T-bet expression and reduced TCR-signaling, resulting in less severe GVHD. Our data demonstrated that *TCF-7*-deficient CD8 T cells utilized NKG2D receptors to kill tumor targets. These findings will have a considerable impact on developing strategies to uncouple GVHD from GVL, and for developing therapeutic interventions for T cell-driven autoimmune disorders.

## Results

### Loss of *TCF-7* in donor CD8 T cells reduced severity and persistence of GVHD symptoms, increased survival from lethal GVHD, and retained anti-tumor capabilities for the GVL effect

Most of the previous research on *TCF-7* utilized a global *TCF-7* knockout because the primary focus was on *TCF-7’s* role as a developmental factor (Gounari and Khazaie, 2022; Weber *et al*., 2011). However, we wanted to study the role of *TCF-7* in mature T cells. Global loss of *TCF-7* results in minimal T cell production from the thymus, because *TCF-7* is critical for DN stages of development (Johnson et al., 2018). Therefore, we obtained mice with a T cell- specific knockout for *TCF-7* (Tcf7 flox/flox x CD4cre, called *TCF-7* cKO here (Xing *et al*., 2016). This allowed us to study mature T cells that developed normally in the thymus, then lost expression of *TCF-7* at the DP phase (Berga-Bolanos et al., 2015).

To determine whether *TCF-7* plays a role in mature alloactivated T cell regulation, which is currently unknown, we used a mouse model of MHC-mismatched allo-HSCT leading to GVHD and GVL. Briefly, BALB/c mice (MHC haplotype d) were lethally irradiated and transplanted with wild-type (WT) bone marrow and C57Bl/6-background (MHC haplotype b) donor CD8 T cells (Mammadli *et al*., 2021c; Mammadli *et al*., 2021d). The donor CD8 T cells came from wild-type (WT), or *TCF-7*-deficient (*TCF-7* cKO) mice. Recipients were given 1X10^6^ CD8 T cells and 10X10^6^ WT T cell-depleted bone marrow cells, as well as 2X10^5^ luciferase-expressing B-cell lymphoma (A-20) cells (Edinger et al., 2003a; Edinger et al., 2003b) to assess GVL responses (Mammadli *et al*., 2021a; Mammadli et al., 2021b; Mammadli *et al*., 2021c; Mammadli *et al*., 2021d). A20 cells are syngeneic to BALB/c mice and allogeneic to C57BL/6 (B6) mice (Edinger *et al*., 2003a; Edinger *et al*., 2003b), meaning that the cells will not be naturally rejected by the BALB/c hosts, but will be attacked by the transplanted donor T cells. The MHC haplotype mismatch between host and donor cells drives alloactivation of donor T cells, which in turn causes GVHD and GVL effects (Hoffmann et al., 2002). To examine disease severity, progression, and recipient mouse survival, recipient mice were weighed and given a clinical score three times per week following transplant, until about day 70 (**Fig. 1A-D**). The mice were scored based on six factors: skin integrity, fur texture, posture, activity level, weight loss, and diarrhea (Cooke et al., 1996). Since the A20 cells express luciferase (called A20 luc) (Mammadli *et al*., 2021c), allowing us to track them by injecting D-luciferin into the recipient mice and imaging them with an *in vivo* bioluminescence scanner (IVIS 50), the mice were scanned one time per week with IVIS 50 until the end of the experiment (**Fig. 1A, 1E**).

**Figure 1:**
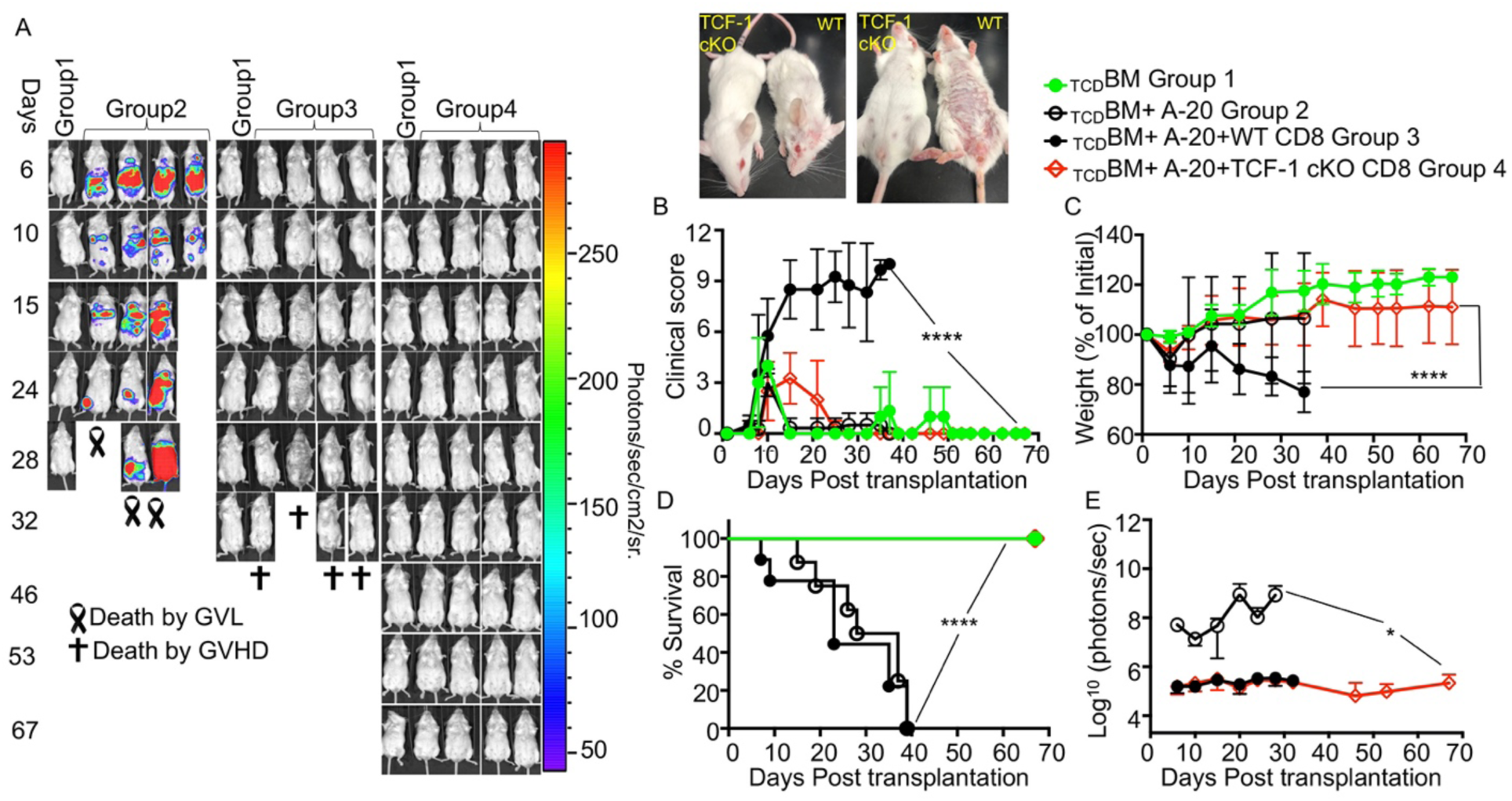
Loss of *TCF-7* in donor CD8 T cells reduces severity and persistence of GVHD symptoms. BALB/c recipient mice (MHC haplotype d) were lethally irradiated and allotransplanted with 1X10^6^ CD8 T cells from WT or *TCF-7* cKO donor mice (MHC haplotype b), as well as 10X10^6^ T cell-depleted bone marrow cells (BM) from WT mice (MHC haplotype b). Recipient mice were also given 2X10^5^ luciferase-expressing A20 tumor cells **(A)** The recipient mice were imaged 1 time a week using IVIS50 for 70 days. Gross pictures of representatives of recipients transplanted with CD8 T cells from *TCF-7* cKO and WT mice at day 25 post-transplant are shown. **(B)** Recipient mice were also given a GVHD clinical score three times per week until day 70, based on combined scores of fur texture, activity level, weight loss, posture, skin integrity, and diarrhea. Mean and SD are plotted, analyzed by one-way ANOVA. **(C)** Weight changes of the recipient mice also were tracked over the time course of disease. **(D)** Survival for each group of recipient mice up to 70 days post-transplant, analyzed by Kaplan-Meier survival analysis. **(E)** Quantification of bioluminescence of tumor growth. For all graphs, * means p-value ≤ 0.05, ** means p-value ≤ 0.01, *** means p-value ≤ 0.001, and **** means p-value ≤ 0.0001 (*n* = 3 mice/group for BM alone: *n* = 4 experimental mice/group for all other groups). Survival is a combination of two experiments. Note: Control mouse is one of the mice from bone marrow only transplanted group used as a negative control for BLI.

We found that mice who received WT donor CD8 T cells had a rapid increase in GVHD severity, with a high score being reached by day 14, suggesting severe GVHD (**Fig. 1B**). This high score was maintained, suggesting persistent disease, and reached a high peak score at day 40 when the recipient mice died of disease burden (**Fig. 1B-D**). WT-transplanted mice lost weight initially and were unable to regain much weight (**Fig. 1C**). In contrast, mice given *TCF-7* cKO CD8 T cells had much better survival, lower peak and average clinical scores, minimal disease burden, and a gain in weight following the initial weight loss period (**Fig. 1B-D**). In addition, the clinical scores for *TCF-7* cKO CD8 T cells transplanted mice quickly reduced to near-control levels following peak score at day 10 (**Fig. 1B**), suggesting that disease does not persist in these mice. Therefore, loss of *TCF-7* in donor T cells led to reduced GVHD severity and persistence, with improved survival (**Fig.1D**).

Regarding anti-tumor effects, we observed that over time, the group receiving only bone marrow and tumor cells showed a large increase in tumor growth (**Fig. 1A, 1E**), because no T cells were present to control the tumor cells. In contrast, most mice given CD8 T cells from any donor type along with the BM and A20 luc cells were able to clear the tumor cells by the end of the experiment (**Fig. 1A, 1E**). The GVL effect was maintained even in *TCF-7* cKO-transplanted mice (**Fig. 1A, 1E**). Altogether, these data show that *TCF-7* is dispensable for GVL effects, but critical for GVHD. Therefore, loss of *TCF-7* in donor T cells provides a clinically optimal phenotype, where GVHD severity is reduced but beneficial GVL effects are maintained.

### Loss of *TCF-7* drives changes to mature CD8 T cell phenotype which are primarily cell- *extrinsic*

It has been shown that loss of *TCF-7* in late stages of T cell development led to impaired output of CD4 T cells, and redirection of CD4 T cells to a CD8 T cell fate(Steinke et al., 2014). To determine whether loss of *TCF-7* affected mature donor T cell phenotype, we performed flow cytometry phenotyping on CD8 T cells (**Fig. 2**). First, we confirmed the loss of *TCF-7* expression in *TCF-7* cKO mice by flow cytometry (**Fig. 2A**). Next, we examined whether the loss of *TCF-7* also altered Eomesodermin (Eomes) and T-box transcription factor 21 (T-bet) expression, both of which are downstream of *TCF-7* (Chen *et al*., 2019). Some reports have claimed that Eomes is activated by *TCF-7* (meaning loss of *TCF-7* reduces Eomes expression) (Paley and Wherry, 2010). However, in our model of conditional *TCF-7* deletion, we found that *TCF-7* cKO CD8 T cells had increased expression of Eomes compared to WT CD8 T cells (**Fig. 2B**). Other reports have claimed that T-bet may be activated or not affected by *TCF-7* (Ma *et al*., 2012), but we found that loss of *TCF-7* led to increased T-bet expression in CD8 T cells (**Fig. 2C**). This suggests that *TCF-7* normally suppresses the expression of Eomes and T-bet in mature CD8 T cells.

**Figure 2.**
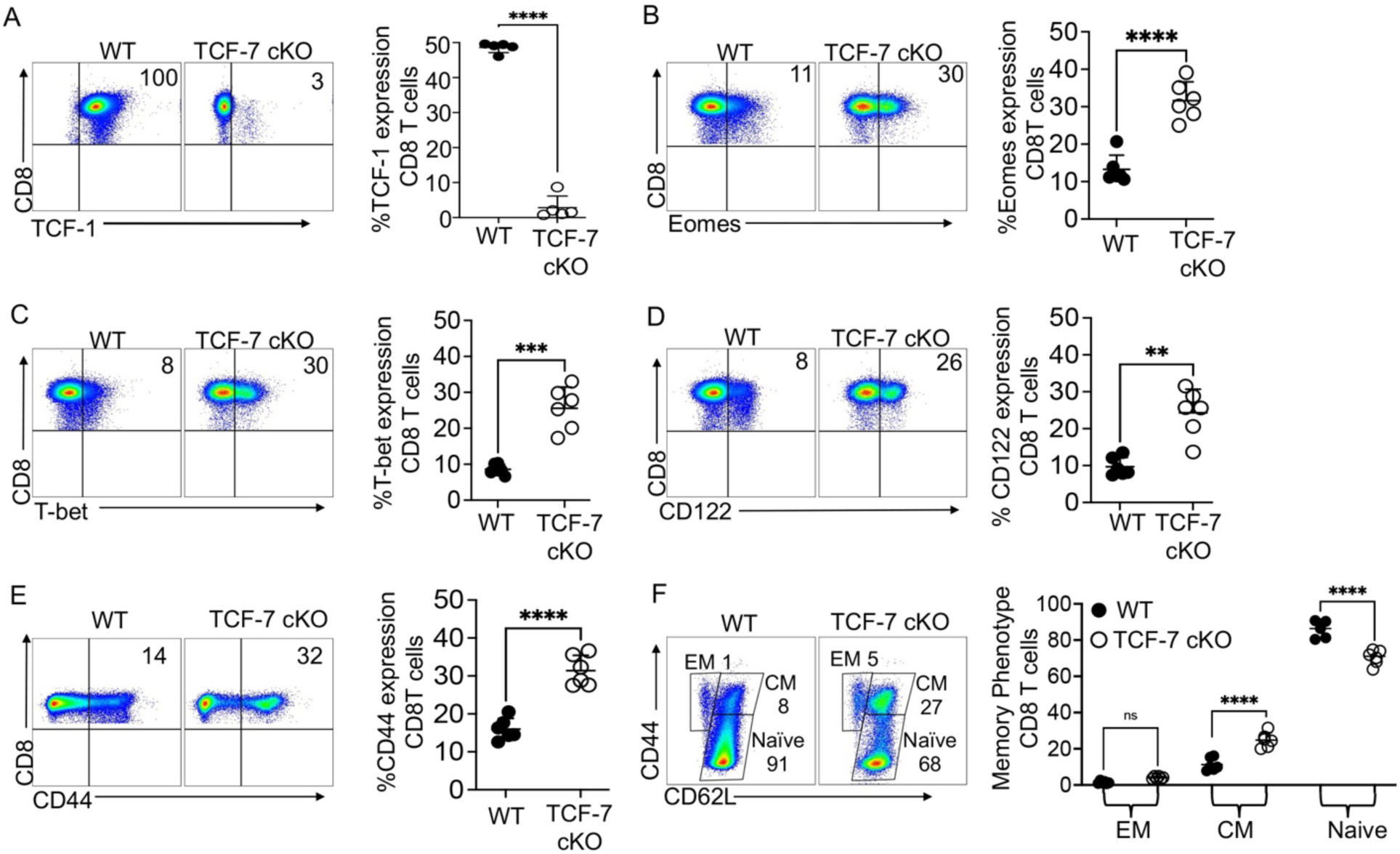
Loss of *TCF-7* changes the mature CD8 T cell phenotype. Naive WT or *TCF-7* cKO donor mice were euthanized and splenocytes were stained for flow cytometry phenotyping**. (A)** Percent of CD8 T cells expressing *TCF-7* and quantified statistical analysis **(B)** Percent of CD8 T cells expressing Eomes and quantified statistical analysis. **(C)** Percent of CD8 T cells expressing T-bet and quantified statistical analysis. **(D)** Percent of CD8 T cells expressing CD122 and quantified statistical analysis. **(E)** Percent of CD8 T cells expressing CD44 and quantified statistical analysis. **(F)** Percent of CD8 T cells expressing central memory, effector memory, or naive phenotypes and quantified statistical analysis. All data are plotted as individual points with mean and SD, all were analyzed with one-way ANOVA, or Student’s t-test (depending on groups). For all graphs, * means p-value ≤ 0.05, ** means p-value ≤ 0.01, *** means p-value ≤ 0.001, and **** means p-value ≤ 0.0001. N=2-3 per group per experiment, with combined data from 3 experiments shown.

Some reports have suggested that CD44^hi^ T cells do not cause GVHD or cause less severe GVHD (Dutt *et al*., 2011). Therefore, we wanted to examine CD8 T cells from *TCF-7* cKO mice for activation markers like CD44 and CD122. Our data showed that CD8 T cells from *TCF-7* cKO mice exhibit increased expression of CD122 and CD44 (**Fig. 2D-E**). Next, using CD62L and CD44 markers, we identified three major memory subsets: central memory (CD44^hi^ CD62L^hi^), effector memory (CD44^hi^ CD62L^low^), and naive (CD44^low^ CD62L ^hi^) cells (**Fig. 2F**). *TCF-7* cKO mice showed increased central memory CD8 T cell subsets and decreased naive CD8 T cells (**Fig. 2F**). Thus, loss of *TCF-7* results in a more memory-skewed phenotype for CD8 T cells. Some reports have suggested that memory T cells delay induction of GVHD (Dutt *et al*., 2011; Mammadli *et al*., 2021c), so this phenotypic change may be beneficial (Nakajima et al., 2021).

Changes to cell phenotype in a knock-out mouse may be cell-intrinsic (due directly to gene deficiency within the cell) or cell-extrinsic (due to changes in the microenvironment from gene deficiency) (Decman et al., 2010; Mammadli *et al*., 2021d). To determine whether the phenotypic effects we observed were cell-intrinsic or cell-extrinsic, we developed a chimeric mouse model. Briefly, we mixed bone marrow from WT and *TCF-7* cKO mice at a 1:4 (WT:TCF) ratio for a total of 50X10^6^ BM cells, then used this mixture to reconstitute lethally irradiated Thy1.1 mice. We used a 1:4 ratio based on our previous published work (Mammadli et al., 2020; Mammadli *et al*., 2021d), to ensure survival of the KO cells (based on our initial observations that *TCF-7* cKO T cells did not proliferate well in culture). At 9 weeks post- transplant, blood was taken to ensure reconstitution and survival of both donor types in each mouse. At 10 weeks, splenocytes were phenotyped by flow cytometry, with donor cells being identified by H2K^b^, CD45.1 (WT), and CD45.2 (*TCF-7* cKO) markers (Mammadli *et al*., 2020).

First, we looked at the *TCF-7* expression in CD45.1+ (WT), and CD45.2+ (*TCF-7* cKO) cells and confirmed that cells from *TCF-7* cKO mice did not express *TCF-7* in chimeric mice **(Supp.Fig.1A).** We did see a statistically significant increase in T-bet expression in CD8 T cells from *TCF-7* cKO donor cells compared to WT donor cells in chimeric mice, when we performed a t-test (data not shown). However, when we compared the T-bet expression in chimeric versus naïve CD8 T cells, we observed that T-bet expression in CD8 T cells from *TCF-7* cKO donor mice was reduced to near-WT levels from elevated levels **(Supp.** **Fig. 1B**). This suggests that the increased expression of T-bet seen in *TCF-7* cKO CD8 T cells from naive mice is a cell-extrinsic effect. Interestingly, in the chimeric mice we observed that Eomes and CD122 expression levels in WT CD8 T cells were significantly increased to near-*TCF-7* cKO levels, suggesting that the increase in Eomes and CD122 expression in CD8 T cells from *TCF-7* cKO mice is primarily cell-intrinsic **(Supp.Fig.1C-D).**

Next, we examined the expression of CD44 and central memory phenotype in chimeric mice. We observed that while the frequencies of these subsets were lower in *TCF-7* cKO-derived CD8 T cells compared to WT-derived CD8 T cells in the chimera (opposite of the trend observed in naive mice), this was because the frequencies of CD44 and CM phenotype in WT cells was enhanced to the levels expressed by *TCF-7* cKO cells from naive mice **(Supp. Fig. 1E-F).** These results suggest that the effects of *TCF-7* deficiency on CD44 and CM phenotype expression in naïve mice could be primarily cell-intrinsic, with cell-extrinsic elements as well. Interestingly, effector memory phenotype in the chimeric mice we observed that levels in WT CD8 T cells were significantly increased to near-*TCF-7* cKO levels, suggesting that the increase in effector memory phenotype in CD8 T cells from *TCF-7* cKO mice is primarily cell-intrinsic **(Supp. Fig. 1G).** Finally, the naïve CD8 T cell population in the chimera coming from *TCF-7* cKO bone marrow was significantly increased compared to CD8 T cells from WT bone marrow and compared to naïve *TCF-7* cKO mice (**Supp. Fig. 1H**). This suggests that the effect on naive CD8 T cells in *TCF-7* cKO mice could be either cell-intrinsic or cell-extrinsic. Altogether, these data suggest that the phenotypic changes seen in *TCF-7* cKO may be primarily cell-intrinsic, with some additional cell-extrinsic effects being present.

### Loss of *TCF-7* alters cytotoxic mediator production in mature CD8 T cells

Our data demonstrated that the loss of *TCF-7* increases Eomes and T-bet expression in mature CD8 T cells (**Fig. 2B-C**). Considering that Eomes and T-bet have been reported to play a central role in anti-tumor responses, we hypothesized that by upregulating Eomes and T-bet, CD8 T cells lacking *TCF-7* can maintain cytotoxicity, and that *TCF-7* is not required for CD8 T cell-mediated cytolytic function (Zhu et al., 2010). We anticipated that CD8 T cells from *TCF-7* cKO mice may have attenuated TCR signaling, so we examined this and other activating receptors by flow cytometry. It is also known that Eomes and T-bet overexpression increases NKG2D expression in NK cells (Kiekens et al., 2021). Considering that loss of *TCF-7* in mature T cells led to upregulation of Eomes and T-bet expression, we hypothesized that loss of *TCF-7* may also lead to upregulation of NKG2D expression in CD8 T cells and enhance the anti-tumor response.

Natural killer group 2 member D (NKG2D) is constitutively expressed on mouse NK cells, NKT cells and some other cells (Abel et al., 2018; Al Dulaimi et al., 2018), but does not get expressed on naïve mouse CD8 T cells (Prajapati et al., 2018). Human CD8 T cells always express NKG2D on their surface, but mouse CD8 T cells only express it upon activation (Wensveen *et al*., 2018) NKG2D is activated by NKG2D ligands (Raulet et al., 2013), and NKG2D ligands are relatively restricted to malignant or transformed cells (Raulet, 2003; Raulet *et al*., 2013). In order to determine whether loss of *TCF-7* affects NKG2D expression and anti-tumor responses, we analyzed NKG2D expression in CD8 T cells. We measured NKG2D expression by flow cytometry before and at different time points after CD3/CD28 activation (Karimi et al., 2015). We found that CD8 T cells from *TCF-7* cKO mice had significantly increased expression of NKG2D on the cells surface compared to CD8 T cells from WT mice, before stimulation (**Fig. 3A**). Next, we wanted to examine whether NKG2D expression was further upregulated on CD8 T cells from *TCF-7* cKO mice compared to CD8 T cells from WT mice after stimulation. CD8 T cells were cultured with 2.5ug/ml anti-CD3 and 2.5ug/ml anti- CD28 antibodies for 24, 48, or 72 hours. These cultured cells were examined for NKG2D expression by flow cytometry. We observed an increase in NKG2D expression on CD8 T cells from both WT and *TCF-7* cKO mice in a time-dependent manner, and at all time points, expression of NKG2D was higher for cells from *TCF-7* cKO mice (**Fig. 3B**). There was no difference in the viability of the cells or CD8 T cell numbers before or after the culture **(Supp.Fig.2A-B).**

**Figure 3:**
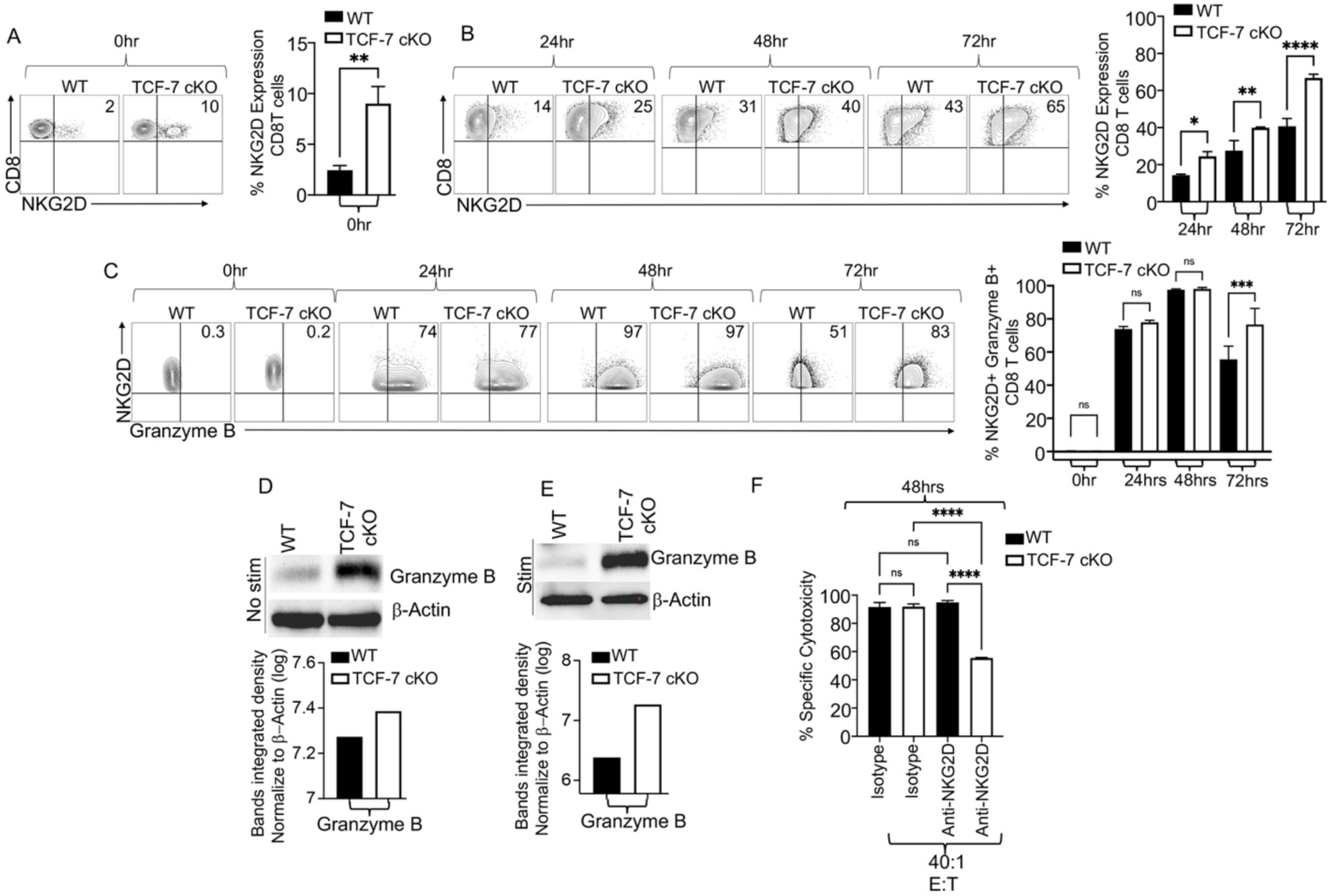
Loss of *TCF-7* alters cytotoxic mediator production by mature alloactivated CD8 T cells. Total splenocytes were isolated from *TCF-7* cKO or WT mice and either left unstimulated or stimulated with anti-CD3/CD28 for 24, 48, or 72 hours in culture. GolgiPlug (1:1000) was added to stimulated samples in each time point except 0-hour samples, and samples were incubated at 37 C with 7% CO2. After 6 hours of culture, the cells were stained with LIVE/DEAD Aqua, and for CD3, CD8, NKG2D, and Granzyme B expression, as determined by flow cytometry. **(A)** Percent of NKG2D expression in freshly isolated CD8 T cells from WT or *TCF-7* cKO mice and quantified statistical analysis. **(B)** Percent of NKG2D expression in 24, 48, or 72 hour-stimulated CD8 T cells from WT and *TCF-7* cKO mice and quantified statistical analysis. **(C)** Percent of Granzyme B expression in unstimulated or 24, 48, or 72 hour-stimulated CD3+ CD8+ NKG2D+ T cells from WT and *TCF-7* cKO mice, and quantified statistical analysis. **(D)** Granzyme B expression in unstimulated CD8 T cells from WT and *TCF-7* cKO mice by Western blot, and bands’ integrated density normalized to β-actin (quantified). **(E)** Granzyme B expression in 10 minute-anti-CD3/CD28-stimulated CD8 T cells from WT and *TCF-7* cKO mice by Western blot, and bands’ integrated density normalized to β-actin (quantified) **(F)** To assess the NKG2D-mediated cytotoxicity, we used luciferase-expressing A20 cells as target cells. Effector cells (MACS-sorted CD8 T cells from *TCF-7* cKO or WT mice) were incubated in 2.5μg/ml anti-CD3 and anti-CD28 coated plates for 48 hours to induce optimal NKG2D expression. Then effector cells were added at 40:1 effector-to-target ratios and incubated at 37°C for 4 hours with A20 cells. Anti-NKG2D antibody (10 μg/mL) or rat IgG1 isotype control antibody (10 μg/mL) was added and incubated for 30 minutes before washing and plating. Triplicate wells were averaged and percent lysis was calculated from the data using the following equation: % specific lysis = 100 × (spontaneous death bioluminescence – test bioluminescence)/ (spontaneous death bioluminescence – maximal killing bioluminescence)(Karimi et al., 2014a; Karimi et al., 2015; Karimi et al., 2014b). N=4 per group with one rappresentative of 2 experiments shown. Western blot were repeated twice and representative examples are shown. All data are shown as individual points with mean and SD, and were analyzed with Student’s t-test or one-way ANOVA (depending on groups). * means p- value ≤ 0.05, ** means p-value ≤ 0.01, and *** means p-value ≤ 0.001.

We also wanted to compare the Granzyme B expression in CD8+, NKG2D+ T cells from *TCF-7* cKO and WT mice (Chu et al., 2013). We did not observe any Granzyme B expression in CD8 T cells before stimulation (**Fig. 3C**). Only 24 hours after stimulation, we observed Granzyme B expression in T cells from both strains, peaking at 48 hours post-stimulation with no difference between strains of mice (**Fig. 3C**). After 72 hours post-stimulation, CD8 T cells from WT mice had significantly reduced Granzyme B expression compared to *TCF-7* cKO CD8 T cells (**Fig. 3C**). We also confirmed total Granzyme B expression in CD8 T cells from *TCF-7* cKO mice, in the presence and absence of CD3/CD28 stimulation, using Western blotting. Total Granzyme B expression was upregulated in CD8 T cells from *TCF-7* cKO mice compared to WT mice **(Fig. 3D-E).** These data demonstrated that CD8 T cells from *TCF-7* cKO mice may maintain anti-tumor responses by killing the target cells with an NKG2D-mediated mechanism, and by persistent upregulation of Granzyme B expression (Liu et al., 2022; Wang et al., 2022).

Next, we wanted to examine the functional consequences of upregulation of NKG2D expression on CD8 T cells from *TCF-7* cKO mice. We utilized an in *vitro* cytotoxicity assay, where we used anti-NKG2D neutralizing antibody. We isolated CD8 T cells from WT and *TCF- 7* cKO mice and cultured them for 48 hours with anti-CD3/anti-CD28 antibodies in order to induce optimal NKG2D expression in CD8 T cells. CD8 T cells from *TCF-7* cKO and WT mice were then cultured with tumor target A20 cells (Edinger *et al*., 2003b) in a 40:1 ratio of tumor cells to CD8 T cells, along with anti-NKG2D antibody or isotype control antibody for 4 hours. We used the A20 cell line as a tumor target because it is known for expressing NKG2D ligands including Rae1, H60, and MULT1(Karimi *et al*., 2015; Nishimura et al., 2008). Triplicate wells were averaged and percent lysis was calculated from the data using the following equation: % specific lysis = 100 × (spontaneous death bioluminescence – test bioluminescence)/ (spontaneous death bioluminescence – maximal killing bioluminescence) (Karimi et al., 2014).

Our data showed that the addition of anti-NKG2D antibody significantly reduced the cytotoxicity of CD8 T cells from *TCF-7* cKO mice, whereas addition of isotype control had no effect on cytotoxicity of the CD8 T cells from *TCF-7* cKO mice (**Fig. 3F**). In contrast, the addition of anti-NKG2D antibody (Karimi *et al*., 2015) did not change cytotoxicity of the CD8 T cells from WT mice (**Fig. 3F**). These data further support the idea that *TCF-7* cKO CD8 T cells maintain their anti-tumor activity through an NKG2D-mediated mechanism. Taking into account that normal tissue does not express NKG2D ligands on the surface and that primarily malignant and transformed cells upregulate these ligands, this could explain why CD8 T cells from *TCF-7* cKO mice cause less GVHD but maintain their anti-tumor activity (Nishimura *et al*., 2008).

### Loss of *TCF-7* alters cytokine production, chemokine expression, and expression of exhaustion markers by mature CD8 T cells

We confirmed that CD8 T cells from *TCF-7* cKO mice mediate cytolytic function primarily through NKG2D. Next, we wanted to examine the mechanism behind why CD8 T cells from *TCF-7* cKO mice induce less GVHD. One of the hallmarks of GVHD is the release of pro-inflammatory cytokines by alloactivated donor T cells, eventually leading to cytokine storm (Lynch Kelly et al., 2015; Mohty et al., 2005). We examined whether loss of *TCF-7* in donor CD8 T cells led to changes in cytokine production, thereby affecting GVHD damage. We allotransplanted lethally irradiated BALB/c mice as described above. Recipient mice were transplanted with 1.5X10^6^ WT or *TCF-7* cKO CD8 donor T cells. Recipients were sacrificed at day 7 post-transplant. Splenocytes were isolated and restimulated by 6 hours of culture with PBS (control) or anti-CD3/anti-CD28 (stimulation), along with Golgiplug. Afterwards, the cultured cells were stained with antibodies against H2K^b^, CD3, CD4, CD8, TNF-α, and IFN-γ. Our data showed that production of TNF-α by donor CD8 T cells trended toward decreasing when *TCF-7* was lost **(Supp. Fig. 3A)**. In contrast, IFN-γ trended toward increasing upon loss of *TCF-7* in CD8 T cells **(Supp. Fig. 3B)**.

We also obtained serum from cardiac blood of recipient mice at day 7 post-transplant and tested it with a mouse Th cytokine ELISA panel (LEGENDplex kit from Biolegend) (Mammadli *et al*., 2020; Mammadli *et al*., 2021d). Levels of TNF-α and IFN-γ in serum of recipient mice given *TCF-7* cKO CD8 T cells were lower than in mice given WT CD8 T cells at day 7 (**Fig. 4A**). In contrast, the serum levels of IL-2 in mice given *TCF-7* cKO CD8 T cells was higher than in mice given WT CD8 T cells at day 7 (**Fig. 4A**). At day 14 post-transplant, the reduction in TNF-α and IFN-γ levels observed at day 7 for *TCF-7* cKO-transplanted mice was preserved (**Fig. 4B**). We observed a trend towards decreased serum levels of IL-2 in mice given *TCF-7* cKO CD8 T cells compared with mice given WT CD8 T cells at day 14 post-transplant, opposite of the effect observed on day 7 post-transplant (**Fig. 4B**). This suggests that *TCF-7* cKO CD8 T cells may be capable of IFN-γ and TNF-α production at the same level as WT CD8 T cells when restimulated, but in reality, produce less IFN-γ and TNF-α than WT cells when allotransplanted.

**Fig. 4.**
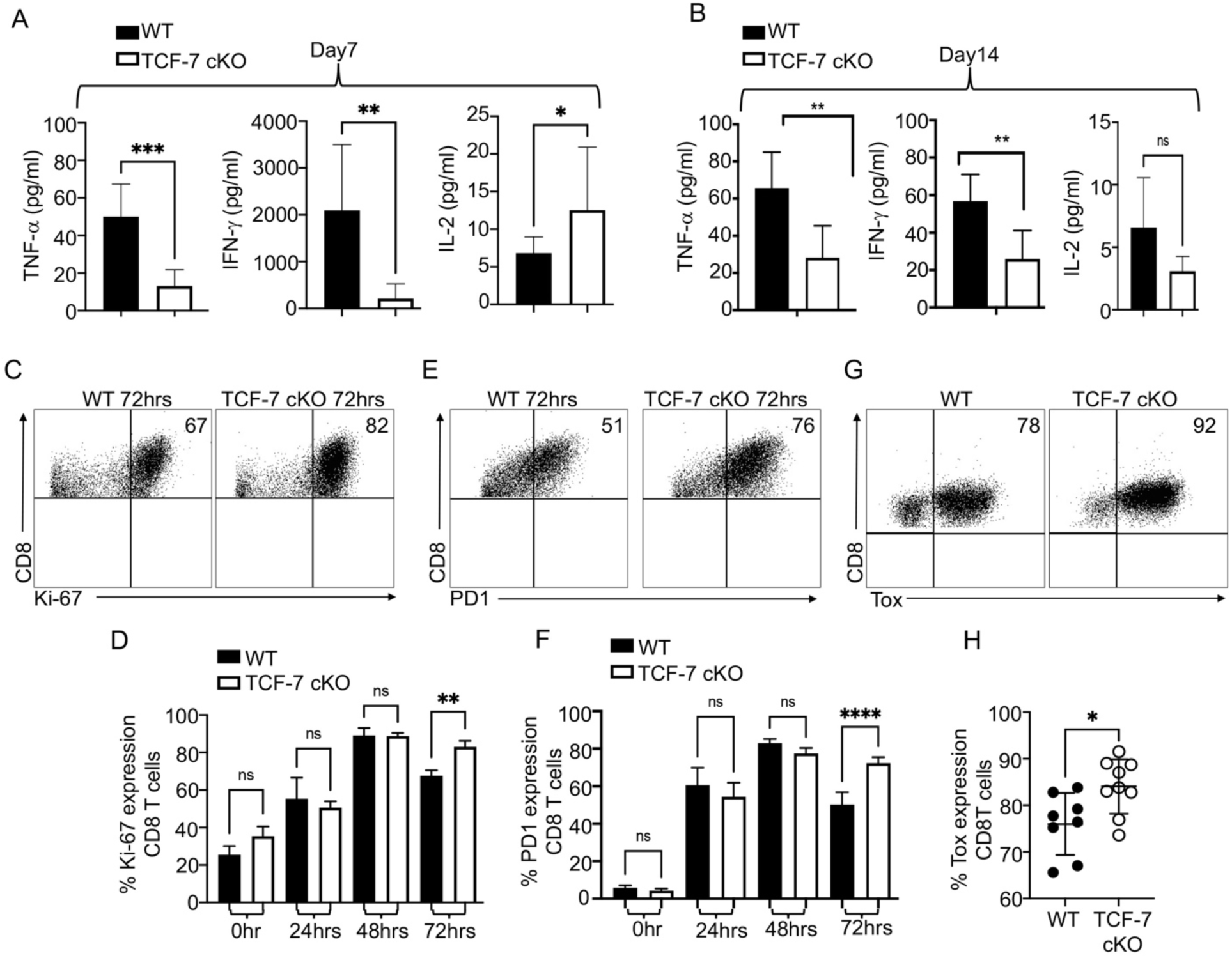
*TCF-7* controls cytokine production and exhaustion of mature alloactivated CD8 T cells. **(A-B)** Recipient mice were allotransplanted with 1.5X10^6^ WT or *TCF-7* cKO donor CD3 T cells as before. At day 7 and day 14 post-transplant, serum was obtained from cardiac blood of allotransplanted recipient mice and tested for various cytokines using a LEGENDplex ELISA assay. **(A)** Serum levels (pg/mL) of TNF-α, IFN-γ, and IL-2 for WT and *TCF-7* cKO- transplanted mice at day 7 post-transplant. **(B)** Serum levels (pg/mL) of TNF-α, IFN-γ, and IL-2 for WT and *TCF-7* cKO-transplanted mice at day 14 post-transplant. **(C-F)** Splenocytes from *TCF-7* cKO and WT mice were obtained and stimulated with anti-CD3/CD28 antibodies for 24hrs, 48hrs, or 72hrs in culture and stained for Ki-67 and PD-1 or were stained immediately after isolation without stimulation. **(C)** Percent expression of Ki-67 in CD8 T cells after 72 hours of anti-CD3/CD28 stimulation in culture determined by flow cytometry. **(D)** Quantification of the *in vitro* Ki-67 expression of CD8 T cells at different time points. **(E)** Percent expression of PD-1 in CD8 T cells after 72 hours of anti-CD3/CD28 stimulation in culture determined by flow cytometry. **(F)** Quantification of the *in vitro* PD-1 expression of CD8 T cells are different time points**. (G-H)** Balb/c mice were allotransplanted as before, with WT or *TCF-7* cKO CD8 donor T cells. On day 7, spleens were removed from recipients, processed to isolate lymphocytes, and spleen-derived lymphocytes from allotransplanted recipient mice were stained for CD3, CD8, H2K^b^, and TOX to identify exhausted T cells. **(G)** Percent expression of TOX in CD8 T cells after allotransplantation (day 7 post-transplant) as determined by flow cytometry. **(H)** Quantification of the TOX expression in *in vivo* donor CD8 T cells. N=3-5 per group for **A-B** with two experiments shown. N=4 per group for **C-F,** one representative of two experiments shown. N=3-5 per group with one representative experiment shown for **G-H**. All data are shown as individual points with mean and SD, and were analyzed with Student’s t-test or two-way ANOVA (depending on groups). * means p-value ≤ 0.05, ** means p-value ≤ 0.01, and *** means p-value ≤ 0.001.

In order for GVHD to persist, donor T cells must proliferate in secondary lymphoid organs and target organs (Beilhack et al., 2005; Ferrara, 2014). Naive and effector T cells drive GVHD, but they are short-lived and must be replaced to maintain an alloresponse (Jiang et al., 2021; Jiang et al., 2014). Also, given that memory cells are increased among CD8 T cells when *TCF-7* is lost, we hypothesized that activation and/or exhaustion of these cells may also be affected. Ki-67 (Blessin et al., 2021) is a marker of T cell activation and proliferation, and TOX (Khan et al., 2019) and PD-1 are markers of exhaustion (Ahn et al., 2018). Therefore, we wanted to determine the Ki-67, TOX, and PD-1 expression levels on WT and *TCF-7* cKO CD8 T cells *in vitro*. We cultured splenocytes from either WT mice or *TCF-7* cKO mice *in vitro* with anti-CD3 and anti-CD28 antibodies for 24, 48, and 72 hours. We did not observe any difference in Ki-67 expression in cells that were not stimulated, but the CD8 T cells from *TCF-7* cKO mice that were stimulated for 72 hours *in vitro* showed significant upregulation of Ki-67 expression (**Fig. 4** **C-D**), suggesting that CD8 T cells from *TCF-7* cKO mice could potentially proliferate more than CD8 T cells from WT mice when restimulated. We also observed the same trend of increased expression for PD-1 at 72 hours post-stimulation (**Fig. 4E-F**). There were no differences in expression of TOX at any time points *in vitro* when CD8 T cells from *TCF-7* cKO mice were compared to WT **(Supp.Fig. 3C).**

Next, we checked the expression of these markers *in vivo* from donor cells that were allo- transplanted in recipient Balb/c mice as described earlier. At day 7 post-transplant, splenocytes were isolated and Ki-67, TOX, and PD-1 expression were detected by flow cytometry. We observed that donor CD8+T cells from *TCF-7* cKO mice expressed more TOX compared to WT, suggesting that donor T cells *TCF-7* cKO mice were more exhausted following allotransplantation (**Fig. 5G-H**). We did not observe any statistically significant differences in Ki- 67 or PD-1 expression at day 7 post-transplant **(Supp.Fig.3D-E).** Taken together these data suggest that CD8 T cells from *TCF-7* cKO mice could be more exhausted than WT CD8 T cells both *in vivo* and *in vitro*.

**Fig. 5.**
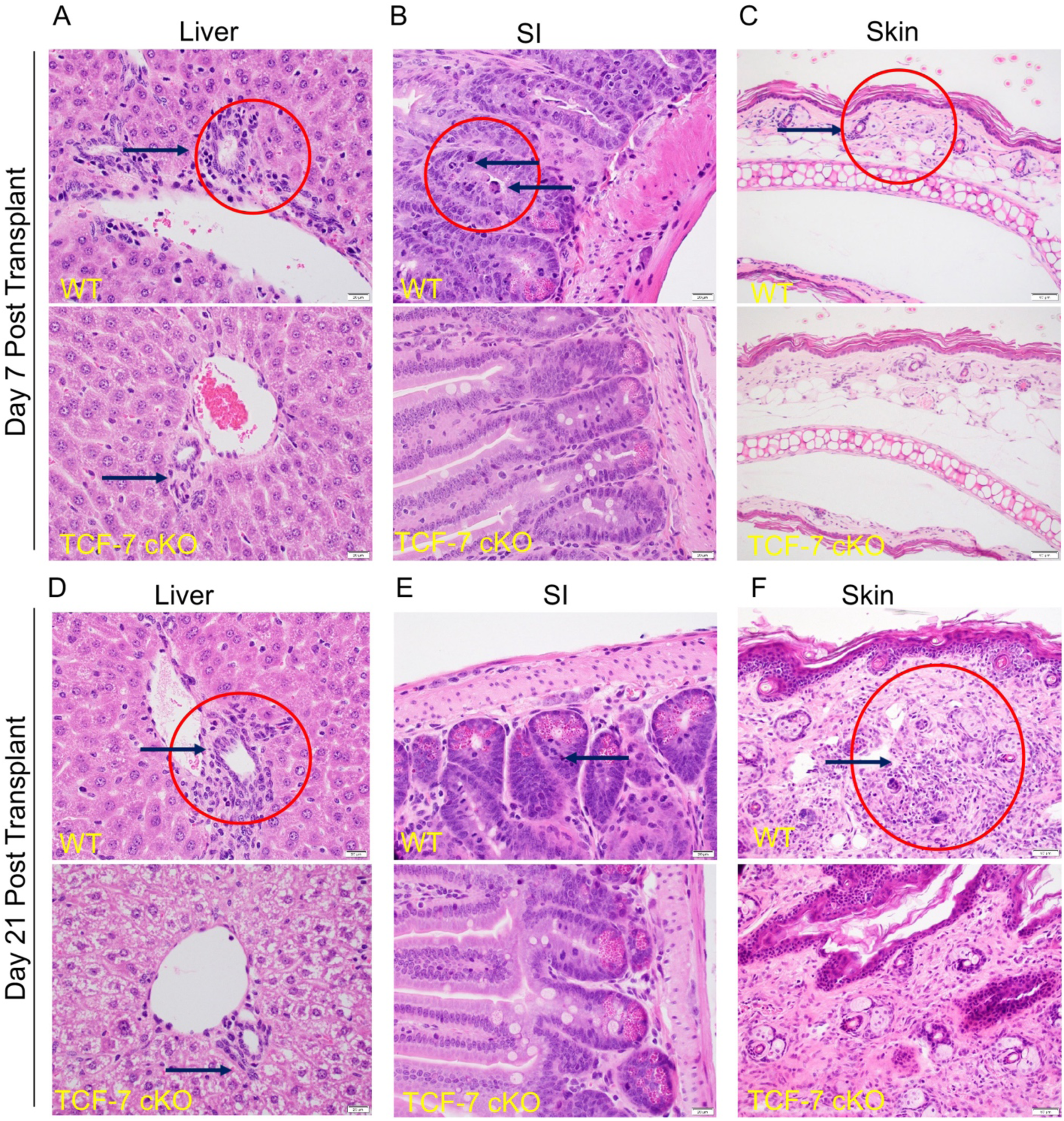
Loss of *TCF-7* in donor CD8 T cells decrease the damage to the GVHD target organs. We collected organs from mice allotransplanted as described above. At day 7 and day 21 post- transplant, organs were taken from recipient mice for histology analyses. Skin, liver, spleen, and small intestine were sectioned, stained with H&E and analyzed by pathologist. Representative sections for each organ per group and timepoint are shown. **(A)** H&E staining of the liver of the recipient mice at day 7 post-transplant (black arrows showing the interlobular bile ducts and red circle showing inflammatory infiltrates). **(B)** H&E staining of the small intestines of the recipient mice at day 7 post-transplant (black arrows showing the crypts of the small intestine and red circle showing apoptotic bodies). **(C)** H&E staining of the skin of the recipient mice at day 7 post-transplant (black arrows showing the dermis of the skin and red circle showing increase of inflammatory cells) **(D)** H&E staining of the liver of the recipient mice at day 21 post-transplant (black arrows showing the interlobular bile ducts and red circle showing inflammatory infiltrates). **(E)** H&E staining of the small intestines of the recipient mice at day 21 post- transplant (blacks arrows showing apoptotic bodies). **(F)** H&E staining of the skin of the recipient mice at day 21 (black arrows showing dermis of skin and red circle showing increased inflammatory cells with destructed adnexal glands).

One of the major functions of alloactivated T cells is migration from spleen to GVHD target organs, including liver and small intestine(Beilhack *et al*., 2005; Ferrara, 2014). Expression of chemokines and chemokine receptors is a critical aspect of T cell migration to target organs. To determine whether expression of these molecules was affected by loss of *TCF- 7* in CD8 T cells, we FACS sorted pre- and post-transplanted donor CD8 T cells from WT or *TCF-7* cKO mice (spleen only for pre-transplant, spleen, or liver for post-transplant). We then extracted RNA from the cells, converted it to cDNA, and performed qPCR using a 96-well mouse chemokine/chemokine receptor plate (Thermo Fisher). As expected, expression of these markers was generally upregulated in alloactivated cells. Expression of these markers was generally higher in *TCF-7* cKO CD8 T cells from pre-transplant spleen compared to WT pre-transplant spleen **(Supp. Fig. 4A)**, but these markers were generally downregulated in *TCF-7* cKO CD8 T cells from post-transplant liver and spleen compared to WT cells **(Supp. Fig. 4B- C)**. Therefore, *TCF-7* controls expression of CD8 T cell chemokine/chemokine receptors.

### Loss of *TCF-7* in donor CD8 T cells led to decreased damage to the GVHD target organs

During GVHD, host tissues are damaged by the activity of alloactivated T cells. To determine whether damage to target organs of GVHD (skin, liver, and small intestine) was altered by loss of *TCF-7* in donor T cells, we collected organs from mice allotransplanted as described above (Beilhack *et al*., 2005; Mammadli *et al*., 2021a; Mammadli *et al*., 2021d; Weeks *et al*., 2021). At day 7 and day 21 post-transplant, we collected pieces of skin, small intestine, and liver from recipient BALB/c mice. These organs were fixed, sectioned, stained with hematoxylin and eosin (H&E), and analyzed by a pathologist (L.S.) (**Fig. 5**). At day 7, *TCF-7* cKO mice showed significantly less inflammatory infiltrates in all the organs. We observed much less inflammatory infiltrates in the bile duct epithelium of the portal triad (black arrows showing the interlobular bile ducts) in the liver of the *TCF-7* cKO-transplanted recipients compared with WT-transplanted recipients (**Fig. 5A**). In the small intestines, no apoptotic bodies were seen in the crypts of the small intestine in the *TCF-7* cKO CD8 T cell-transplanted mice, while frequent apoptotic bodies were present in the WT CD8 T cell-transplanted mice at day 7 post-transplantation (black arrows) (**Fig. 5B**). In the skin, a mild increase in inflammatory cells was observed in the dermis of the WT CD8 T transplanted mice, while the dermis of the *TCF-7* cKO CD8 T transplanted mice appears normal at day 7 post-transplantation (**Fig. 5C**).

Again, at day 21 post-transplant, *TCF-7* cKO CD8 T cell transplanted mice showed significant less inflammatory infiltrates in all the sectioned GVHD target organs. We observed much less inflammatory infiltrates involving the bile duct epithelium of the portal triad (black arrows showing the interlobular bile ducts) in the liver of the *TCF-7* cKO CD8 T cell-transplanted mice compared with WT CD8 T cell-transplanted mice (**Fig. 5D**). At day 21 post-transplant, no apoptotic bodies were seen in the crypts of the small intestines of the *TCF-7* cKO CD8 T cell transplanted mice, while few apoptotic bodies are present in the small intestines of the WT CD8 T cell transplanted mice (black arrows) (**Fig. 5E**). A marked increase in inflammatory cells (red circle) and destruction of the adnexal glands was observed in the dermis of the WT CD8 T cell-transplanted recipients, while the dermis of the *TCF-7* cKO CD8 T cell transplanted recipients showed only a mild increase in dermal inflammatory cells, with preservation of adnexal glands (**Fig. 5F**). Altogether, these data suggest that *TCF-7* normally promotes GVHD damage to healthy tissues and is indispensable for T cell-driven damage. Thus, loss of *TCF-7* in donor T cells leads to reduced severity and persistence of GVHD over time.

### *TCF-7* alters the transcriptomic signature of alloactivated T cells

Given that the phenotype and functions of donor T cells, as well as disease outcomes, were significantly altered by loss of *TCF-7* on donor cells, we sought to determine what specific gene changes occurred to support this. We allotransplanted recipient BALB/c mice with 1X10^6^ donor CD3 T cells as above, and FACS-sorted pre- and post-transplant WT or *TCF-7* cKO donor CD8 T cells, which were stored in Trizol and transcriptionally profiled. When we analyzed the genetic profile of the pre- transplanted CD8 T cells from *TCF-7* cKO and WT mice, we were not able to determine any differentially expressed genes (DEGs, FDR<0.1). However, when we performed Gene Set Enrichment analysis (GSEA) using the Hallmark pathways collection from Molecular Signatures Database (MSigDB), we identified that cytokine signaling pathways like TNF-a Signaling via NF-κβ and Interferon gamma response were enriched in pre-transplanted CD8 T cells from WT mice compared with cells from *TCF-7* cKO mice **(Supp.Fig.5A-C)**. Meanwhile, a number of pathways involved in cell cycle also were enriched in WT cells versus *TCF-7* cKO pre- transplanted CD8 T cells, like the P53 pathway, G2M checkpoint, DNA repair, and Myc targets pathways **(Supp.Fig.5A).** MTOR signaling, Allograft rejection, and Oxidative phosphorylation pathways were also enriched in pre-transplant CD8 T cells from WT mice **(Supp.Fig.5A,6D-F),** suggesting that loss of *TCF-7* alters the transcriptional profile of CD8 T cell towards decreased cytokine release while also altering the cell cycle, leading to a lessening of the alloactivation responses.

When we analyzed the post-transplanted CD8 T cells which were alloactivated *in vivo*, we identified 2548 differentially expressed genes (DEGs; FDR<0.1) when comparing *TCF-7* cKO cells to WT cells (**Fig. 6A-B**). A majority of the DEGs (2000 genes) were downregulated (module 2 in heatmap) and only 548 genes (module 1 in heatmap) were upregulated in post-transplant CD8 T cells from *TCF-7* cKO mice compared to WT mice (**Fig. 6A-B**). We analyzed both up- and downregulated DEGs for the Gene Ontology (GO) enrichment analysis using Functional Annotation Chart tools, selecting only the top 20 GO BP (Biological Process) terms in the Database for Visualization and Integrative Discovery (DAVID)(Huang da et al., 2009; Sherman et al., 2022). The analysis showed that the differentially expressed genes in post-transplant CD8 T cells from *TCF-7* cKO mice compared to WT were involved in Cell Cycle, Cell-cell adhesion, Cell division, Apoptotic process, Antigen processing and presentation via MHC class I, TCR signaling, Regulation of NF-κB signaling, Metabolic process, and others **(Supp.Fig.6).** Once we knew which processes these DEGs played a role in, we pulled out the top genes that were altered for each GO BP term that we were interested in. When we looked at the top 35 DEGs based on P-value that were altered in Cell cycle, we observed that a majority of them were downregulated in *TCF-7* cKO compared to WT (**Fig. 6C**). We also looked at the top 30 genes based on P-value that were altered in Apoptotic processes (**Fig. 6D**), which also showed that most of the genes were downregulated in CD8 T cells in *TCF-7* cKO mice. When we looked at the top 30 genes that play a role in metabolic processes, we observed that only 1 gene was upregulated, and the rest were downregulated in *in vivo* alloactivated CD8 T cells from *TCF-7* cKO mice (**Fig. 6E**). While the upregulated genes in the NF-κB pathway were Fasl, Ubd, Chuk and others, the downregulated genes were Rela, Irf3, Traf2, Mavs, Tradd, Nod1, Tnfrsf1a, and Trim25 (**Fig. 6F**).

**Fig. 6.**
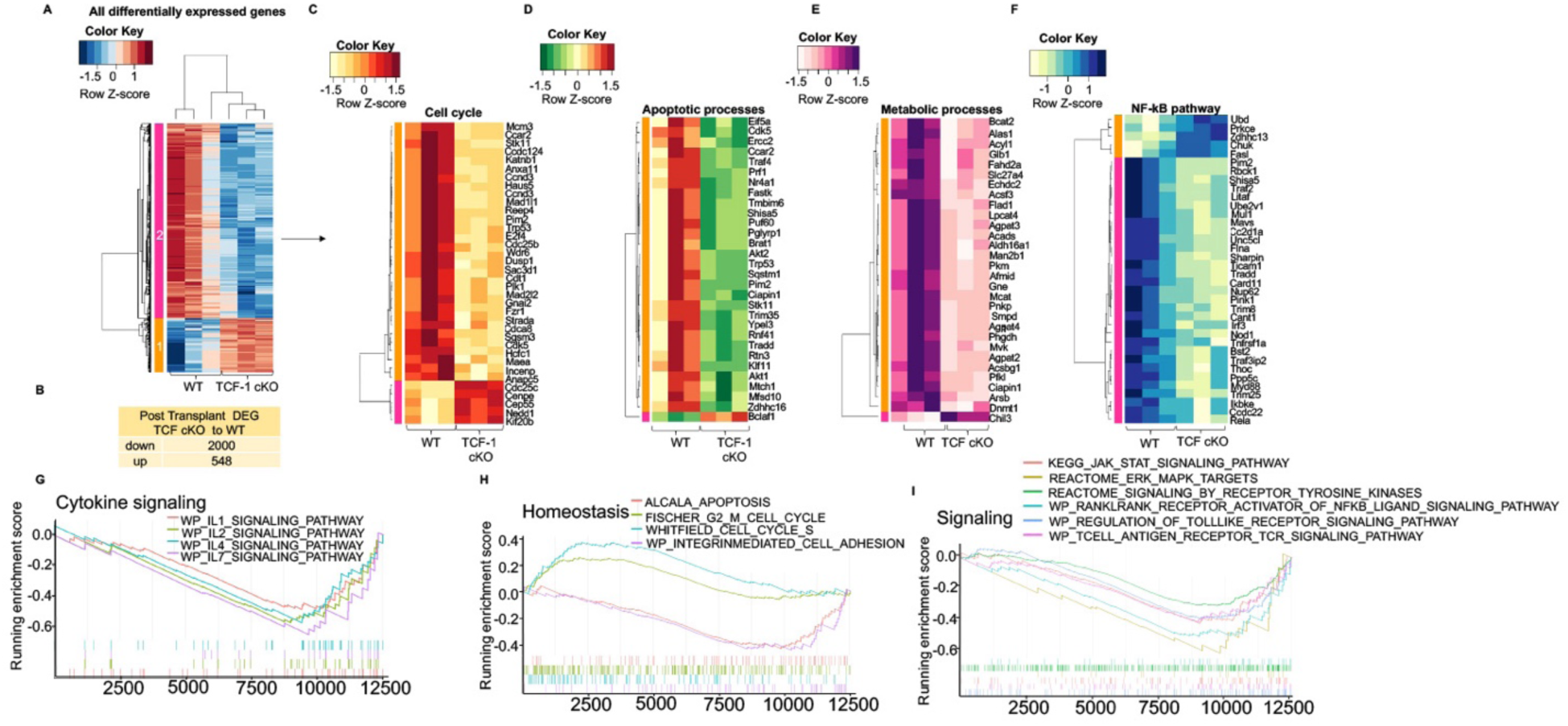
Loss of *TCF-7* changes the genetic signature of alloactivated donor CD8 T cells. CD8 T cells from WT and *TCF-7* cKO mice were FACS sorted either into Trizol as pre- transplanted samples, or into 10% FBS-containing media for transplantation into recipient mice as described previously. At day 7 post-transplant, donor CD8 T cells were FACS-sorted back from recipient spleens of *TCF-7* cKO- and WT-transplanted mice. RNA was extracted and prepped by the Molecular Analysis Core (SUNY Upstate). Paired end sequencing was done with an Illumina NovaSeq 6000 system at the University at Buffalo Genomics Core. For data analysis, we used the statistical computing environment R (v4.0.4), the Bioconductor suite of packages for R, and RStudio (v1.4.1106). We calculated the transcript abundance by performing pseudoalignment using Kallisto. **(A)** Hierarchical clustering of genes and samples, heatmap illustrating the expression of the differentially expressing genes (DEG’s; FDR<0.1) of post- transplanted CD8 T cells *TCF-7* cKO compared to WT. **(B)** Table showing the number of up- or downregulated DEGs in post-transplanted CD8 T cells (*TCF-7* cKO compared to WT). **(C)** Heatmap showing top significant 35 differentially expressed genes that play a role in the Cell Cycle pathway, identified using DAVID Functional Annotation Analysis and GO-BP terms in post-transplanted CD8 T cells (*TCF-7* cKO compared to WT)**. (D)** Heatmap showing top significant 30 differentially expressed genes that play a role in the Apoptotic Processes pathway, identified using DAVID Functional Annotation Analysis and GO-BP terms in post-transplanted CD8 T cells (*TCF-7* cKO compared to WT). **(E)** Heatmap showing top significant 30 differentially expressed genes that play a role in the Metabolic Processes pathway, identified using DAVID Functional Annotation Analysis and GO-BP terms in post-transplanted CD8 T cells (*TCF-7* cKO compared to WT). **(F)** Heatmap showing top significant 35 differentially expressed genes that play a role in the NF-κB pathway, identified using DAVID Functional Annotation Analysis and GO-BP terms in post-transplanted CD8 T cells (*TCF-7* cKO compared to WT). **(G)** Gene set enrichment analysis (GSEA) enrichment plots of Cytokine signaling pathways, including IL-1, IL-2, IL-4, and IL-7 signaling pathways from WP terms that are enriched in post-transplanted CD8 T cells from WT mice (versus *TCF-7* cKO). Negative Enrichment Score is an indicator of downregulation, and positive Enrichment Score is an indicator of upregulation of the genes in the post-transplanted CD8 T cells from *TCF-7* cKo mice. DAVID enrichment scores >1.3 are equivalent to a P value<0.05 (**H)** GSEA enrichment plots of G2 to M Cell Cycle and Cell Cycle S pathways that are enriched in *TCF-7* cKO mice, and Integrin Mediated Cell Adhesion and Apoptosis pathways that are enriched in post- transplanted CD8 T cells from WT mice. **(I)** GSEA enrichment plots of JAK-STAT signaling, Toll like receptor signaling, T cell receptor signaling, ERK MAPK signaling, signaling by Tyrosine kinases, Rankl - Rank mediated NF-kB signaling pathways that are enriched in post- transplanted CD8 T cells from WT mice. Again, Negative Enrichment Score is an indicator of downregulation, and positive Enrichment Score is an indicator of upregulation of the genes in the post-transplanted CD8+ T cells from *TCF-7* cKo mice. DAVID enrichment scores >1.3 are equivalent to a P value<0.05.

Gene Set Enrichment analysis (GSEA) using the Hallmark pathways identified that signaling pathways like PI3K-AKT-MTOR Signaling and TNFA Signaling via NF-κβ were enriched in post-transplanted CD8 T cell from WT mice compared to cells from TCF-7 cKO mice **(Supp.Fig.7A-B)**. GSEA analysis using the C2 canonical pathways showed that a number of cytokines signaling pathways involving IL-1, IL-2, IL-4, and IL-7 were enriched in post- transplanted CD8 T cells from WT mice compared to cells from *TCF-7* cKO mice (**Fig. 6G**). While Cell cycle pathways were enriched in CD8 T cells from *TCF-7* cKO mice compared to cells from WT mice, Apoptosis and Cell adhesion pathways were enriched in post-transplanted CD8 T cells from WT mice (**Fig.6H**). Meanwhile, a number of cells signaling pathways were also enriched in WT CD8 T cells compared to *TCF-7* cKO cells, such as TCR signaling, Toll like receptor signaling, Jak-Stat signaling, ERK-MAPK signaling, and NF-kB pathways (**Fig. 6I**).

We also analyzed the genes that were altered in KEGG pathways, which revealed that DEGs that were altered in *in vivo* alloactivated CD8 T cells from *TCF-7* cKO mice (compared to WT) were involved in pathways like Cell cycle, DNA replication, Metabolic pathways, Natural killer mediated cytotoxicity, TCR signaling, JAK-STAT signaling, Chemokine receptor signaling, and others (**Fig. 7A**). Specifically, Klrk1 gene for NKG2D on Natural killer mediated cytotoxicity pathway were enriched in allo-activated *TCF-7* deficient CD8+ T cells compared to WT CD8+ T cells. Clustering of genes that were affected in TCR signaling showed that while AKT1, AKT2, Pik3r5, Zap70, LCK, Lat, PLCγ1, Pdcd1, Vav1, Rela, Mapk3, Nfkbia, and Nfatc1 were downregulated, Ifng and Ptprc were upregulated in post-transplanted CD8 T cells from *TCF-7* cKO mice (**Fig. 7B**). The JAK/STAT signaling pathway is important for cytokine production and for the response of T cells to cytokines. Analysis revealed that IL2RB, JAK3, STAT5B, STAT3, STAT1, Cish, Il2rb, Socs3, and Socs1 genes were downregulated in the JAK-STAT pathway for *TCF-7* cKO CD8 T cells compared to WT (**Fig. 7C**).

**Fig. 7.**
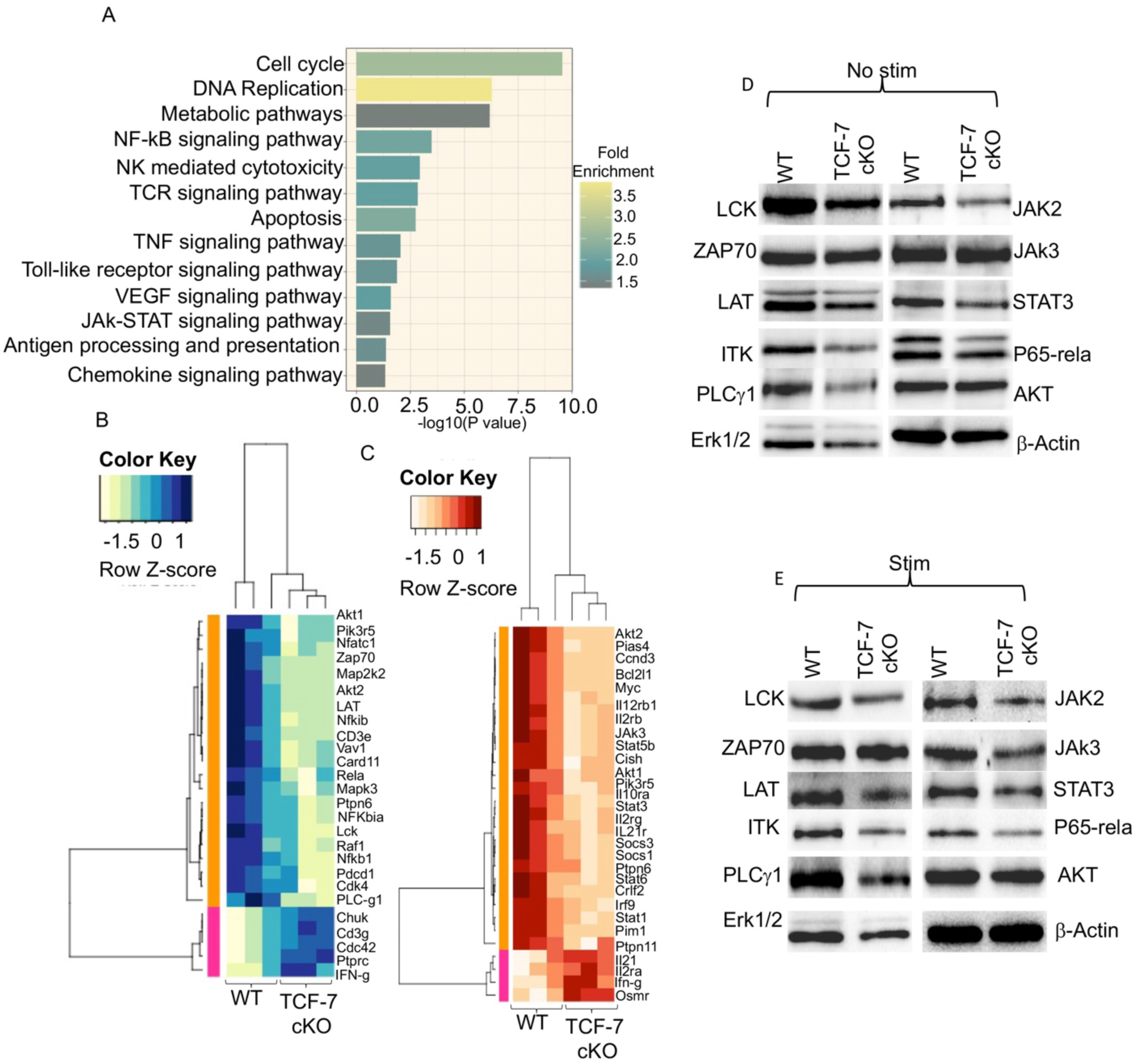
Loss of *TCF-7* decrease TCR, JAK-STAT, and NF-kB signaling downstream. **(A)** A bar plot showing KEGG (Kyoto Encyclopedia of Genes and Genomes) pathways identified by David functional annotation analysis. **(B)** Heatmap showing the altered genes in TCR signaling pathway from KEGG pathways**. (C)** Heatmap showing the altered genes in JAK-STAT signaling pathway from KEGG pathways. **(D)** Western blot showing the protein expression levels of LCK, ZAP70, LAT, ITK, PLCγ1, ERK1/2, JAK2, JAK3, STAT3, P65-RelA, AKT, and β-actin of freshly isolated CD8 T cell lysates from *TCF-7* cKO and WT mice**. (E)** Western blot showing the protein expression levels of LCK, ZAP70, LAT, ITK, PLCγ1, ERK1/2, JAK2, JAK3, STAT3, P65-RelA, AKT, and β-actin in 10 minute-CD3/CD28-stimulated CD8 T cell lysates from *TCF-7* cKO and WT mice. All the western blots repeated at least three and one representative of each protein and quantification is shown.

In order to confirm these changes in genes, we isolated CD8 T cells from WT and *TCF-7 cKO* mice and determined the baseline level protein expression of LCK, ZAP70, LAT, ITK, PLCγ1, ERK1-2, Jak2, Jak3, Stat3, p65-rela, AKT, and actin in unstimulated and 10 minute-anti- CD3/CD28-stimulated samples. Data from non-stimulated samples revealed that protein expression of most of the markers was downregulated in CD8 T cells from *TCF-7* cKO mice compared to WT mice; only ZAP70, JAK3 and AKT were not affected by loss of *TCF-7* **(Fig. 7D, Supp.Fig. 8A).** We observed even more robust differences in samples that were stimulated with anti-CD3/CD28 for 10 minutes, and again only ZAP70 and AKT were unaffected by loss of *TCF-7* **(Fig. 7E, Supp.Fig.8B).** Altogether, the data from RNA sequencing analysis and Western blots of stimulated and unstimulated samples showed attenuation of TCR signaling and many other pathways in CD8 T cells from *TCF-7* cKO mice. These results help to explain why CD8 T cells from *TCF-7* cKO mice cannot induce GVHD as severely as CD8 T cells from WT mice.

## Discussion

T Cell Factor-1 (*TCF-7*) is a critical regulatory transcription factor in T cell development and functions (Zuniga-Pflucker, 2004). *TCF-7* is known to be important for T cell development, as well as activation in some contexts (Yu et al., 2010). *TCF-7* has been extensively studied in viral infection(Escobar *et al*., 2020; He *et al*., 2016; Im *et al*., 2016; Kurtulus *et al*., 2019; Miller *et al*., 2019; Siddiqui *et al*., 2019; Utzschneider *et al*., 2016; Weber *et al*., 2011; Wu *et al*., 2016). However, the role of (*TCF-7*) in a mouse model of allogeneic transplant has not been investigated. Using a murine allogeneic transplant model, we have shown that CD8 T cells from *TCF-7* cKO effectively clear tumor cells without inducing GVHD by producing significantly less inflammatory cytokines (Ju *et al*., 2005; Seif *et al*., 2017). Our data also uncovered that CD8 T cells from *TCF-7* cKO mice cause significantly less tissue damage in GVHD target organs (Bleakley *et al*., 2012; Breems and Lowenberg, 2005). Here, we show that loss of *TCF-7* alters a number of CD8 T cell functions, and while it is dispensable for anti-tumor responses, it is essential for host tissue damage, cytokine production and signaling, and gene expression, playing a role in a number of immunological and biological pathways during alloactivation. (Yu *et al*., 2010). This murine model allows us to study T cell function, clinical outcomes, and gene expression all in one model. Here, we showed that *TCF-7*-deficiency alters the phenotype of CD8 T cells by upregulating CD44 and CD122. We and other have shown that innate memory-like CD8 T cells expressing CD12hi and CD44 hi, Eomes and T-bet ameliorate GVHD development (Huang et al., 2019; Karimi *et al*., 2014; Mammadli *et al*., 2020; Mammadli *et al*., 2021d; Zheng et al., 2009). The role of central (CD44^hi^, CD62L^hi^) and effector (CD44^hi^, CD62L^low^) memory phenotypes has been investigated previously (Huang *et al*., 2019; Zheng *et al*., 2009). Our data demonstrated that *TCF-7* significantly impacts CD8 T cell central memory phenotypes, suggesting that *TCF-7* might be a regulator for T cell activation. While effector and naive cells are known to cause severe GVHD, central memory cells are often associated with less severe disease (Dutt *et al*., 2011; Tugues et al., 2018; Zheng *et al*., 2009). Our findings suggesting that this phenotypic change in *TCF-7 cKO* cells may be beneficial for reducing disease severity. Our experiments in mixed bone marrow chimeras showed that bone marrow-derived CD8 T cells from WT and or *TCF-7* cKO mice developed in the same thymus have a similar phenotype to each other, and a different phenotype than CD8 T cells from naïve WT or *TCF-7* cKO mice. We found that the upregulation of activation marker expression like Eomes, CD122, and the effector memory phenotype is primarily cell-intrinsic, with changes to other markers being either cell-extrinsic or primarily cell-intrinsic with other extrinsic effects.

Donor T cells are crucial for target organ injury in graft-versus-host disease (GVHD). These alloactivated T cells proliferate, migrate to target organs (liver, skin, and small intestine), and produce cytokines during GVHD(Bastien et al., 2012; Reddy and Ferrara, 2008; Villarroel et al., 2014). Our data showed that tissue damage in liver, skin, and small intestine (all target organs) was reduced by loss of *TCF-7* in donor CD8 T cells at all timepoints. This shows that *TCF-7* in donor T cells is required for GVHD damage and persistence over time.

Donor T cells eliminate tumor cells (GVL) but also cause graft-versus-host disease (GVHD) (Bleakley *et al*., 2012; Tugues *et al*., 2018). Our data showed that CD8 T cells from *TCF-7* cKO mice were able to clear tumor without causing GVHD, suggesting that *TCF-7* is dispensable for anti-tumor responses. Our data revealed that CD8 T cells from *TCF-7* cKO mice mediate cytolytic function via NKG2D. We also confirmed this hypothesis by blocking the surface NKG2D by anti-NKG2D antibody on CD8 T cells from *TCF-7* cKO and WT mice and performing an *in vitro* cytotoxicity assay. While the anti-tumor response of TCF1-deficient CD8 T cell against A20 cells (which express the NKG2D ligands like Rae1, H60, and MULT1) (Nishimura *et al*., 2008) was diminished, cytotoxicity of WT CD8 T cells was not affected. Upregulation of the *KLRK1* gene (also known as NKG2D) in alloactivated *TCF-7* deficient CD8+ donor T cells further confirmed our hypothesis. Furthermore, the increase in Granzyme B expression in *TCF-7* cKO CD8 T cells by flow cytometry and Western blot also provides evidence as to why the anti-tumor response is preserved despite weakened TCR signaling in TCF-7 cKO CD8 T cells (Presotto et al., 2017).

Once we had a clear mechanism for the GVL effect, we looked at functions that were altered in CD8 T cells from *TCF-7* cKO mice that could produce the attenuated GVHD effect. We discovered that the serum levels of cytokines like TNFα and IFNγ were decreased in *TCF-7* deficient CD8 T cell-transplanted mice. Published data has shown the CD8 T cell lacking *TCF-7* develop exhaustion while clearing viral infection(Gautam et al., 2019; Seo et al., 2019) While the expression of TOX, an exhaustion marker, was not affected in freshly isolated or *in vitro-* stimulated *TCF-7* deficient CD8 T cells, alloactivated *TCF-7*-deficient CD8 donor T cells upregulated TOX at day 7 post-transplant (Scott et al., 2019). Another exhaustion marker, PD-1, was upregulated after 72 hours in *in vitro-*stimulated CD8 T cells from TCF-7 cKO mice, but *in vivo* expression of PD-1 in donor T cells was not increased (Wang et al., 2019a; Xu et al., 2019). This could be explained by differences between alloactivation *in vivo* and TCR-mediated activation *in vitro.* Ki-67, a proliferation marker, was also altered by loss of *TCF-7*, suggesting that CD8 T cells from *TCF-7* cKO mice may proliferate more compared to WT CD8 T cells (Sobecki et al., 2016). We also confirmed that CD8 T cells from *TCF-7* cKO mice cause less tissue damage to the target organs, and loss of *TCF-7* alters chemokine receptor expression both pre- and post-transplant, which could explain why these cells cause less severe GVHD with increased survival of recipient mice (Ferrara, 2014).

These studies provide evidence for how *TCF-7* regulates the functions of peripheral T cells. The phenotype caused by loss of *TCF-7* is clinically optimal, because it allows for clearance of residual malignant cells while limiting the risk of life-threatening GVHD damage (Guinan et al., 1999). This observation, coupled with the increase in exhaustion of *TCF-7* cKO donor CD8 T cells, suggests that donor cells lacking *TCF-7* are highly activated and cytotoxic to malignant cells early on following transplant, but quickly become exhausted, limiting GVHD progression.

Finally, to identify the changes to the genetic program of donor T cells that occurred in the absence of *TCF-7*, we performed RNA sequencing analysis on pre- and post-transplant donor T cells. Even though we could not identify the Differentially Expressed Genes (DEGs) in pre-transplanted T cells (which we attribute to technical problems), gene set enrichment analysis revealed that loss of *TCF-7* alters the genetic signature of pre-transplanted CD8 T cells. A number of signaling pathways involved in cytokine production and cell cycle were enriched in pre-transplanted CD8 T cell from WT mice compared to cells from *TCF-7* cKO mice, suggesting that loss of *TCF-7* alters the transcriptional profile of the CD8 T cell towards decreased cytokine release, while altering the cell cycle in baseline and leading to the lessening of the allo-activation response. Meanwhile, 2548 DEGs (FDR<0.1) were identified when comparing post-transplanted CD8 T cells from *TCF-7* cKO mice and WT mice. Both GO Annotation analysis of DEGs and Gene Set Enrichment analysis revealed that a number of pathways such as Cell cycle, DNA replication, Metabolic pathways, TCR signaling, JAK-STAT signaling, and Chemokine receptor signaling were enriched in post-transplanted CD8 T cells from WT mice compared with cells from *TCF-7* cKO mice. Transcriptomic analysis also revealed that KLRK-1 gene for NKG2D were upregulated in allo-activated *TCF-7* cKO CD8+ T cells compared to the WT CD8+T cells, which confirming our findings in in-vitro cytotoxicity assay and flow cytometry data. These findings suggest that loss of *TCF-7* leads to changes in the transcriptomic profile of the CD8 T cells towards producing less cytokines and attenuating the T cell response, while increasing cytotoxicity. Chemokine receptor pathways were also enriched in alloactivated WT CD8 T cells which confirmed the qPCR analysis of chemokine receptors. The decrease in chemokine receptors helps to explain why we observed less tissue damage in GVHD target organs after allo- transplantation, and attenuated GVHD persistence, in mice given *TCF-7* cKO CD8 T cells. By using Western blotting, we also confirmed changes in TCR and JAK/STAT signaling before and after stimulation with anti-CD3/CD28, which helps to explain why we saw less serum cytokines at day 7 and day 14 post-transplant in mice given *TCF-7* cKO CD8 T cells. This also helps to explain why GVHD didn’t persist in recipients given CD8 T cells which lack *TCF-7*.

Altogether, these data suggest that *TCF-7* is a major transcription factor that plays a role in T cell development. Our work shows that *TCF-7* is dispensable for cytotoxic function of mature alloactivated CD8 T cells but is indispensable for GVHD. TCR, JAK-STAT, and NF-κB signaling as well as cytokine production, these findings will help to establish an understanding of *TCF-7* as a critical factor in the GVHD/GVL regulatory network of CD8 T cells.

## Materials and Methods

### Mouse Models

For transplant, the following female donor mice were used: B6-Ly5.1 (CD45.1+, “WT” or B6.SJL-Ptprca Pepcb/BoyCrl, 494 from Charles River), C57Bl/6J (CD45.2+, “WT”, 000664 from Jackson Laboratories), or Tcf7 flox x CD4cre (referred to here as “ *TCF-7 cKO*” (Ma *et al*., 2012), obtained from Dr. Jyoti Misra Sen at NIH by permission of Dr. Howard Xue, and bred in-house),. These donor mice were age-matched to each other and to recipients as closely as possible. BALB/c female mice (CR:028 from Charles River, age 6-8 weeks or older) were used as recipient mice for transplant experiments, and Thy1.1 mice (B6.PL-Thy1a/CyJ, 000406 from Jackson Labs) were used as recipient mice for chimera experiments.

### Allotransplant and Tumor Models

BALB/c recipient mice were irradiated with two doses of 400 cGy of x-rays (total dose 800 cGy) and rested for at least 12 hours between doses. Mice were also rested for 4 hours prior to transplantation. T cells (total CD3+ or CD8+) were separated from WT and *TCF-7 cKO* spleens using CD90.2 or CD8 microbeads and LS columns (Miltenyi, CD8: 130-117-044, CD90.2: 130-121-278, LS: 130-042-401). 1X10^6^ donor cells (unless otherwise mentioned) were injected IV into the tail vein in PBS, along with 10X10^6^ WT bone marrow cells. Bone marrow was T-cell depleted with CD90.2 MACS beads (130-121-278 from Miltenyi) and LD columns (130-042-901 from Miltenyi). For short-term experiments, at day 7 post-transplant, recipient mice were euthanized and serum, spleen, lymph nodes, small intestine, or liver were collected, depending on the experiment. For GVHD and GVL experiments, recipient mice were also given 2X10^6^ luciferase expressing B-cell lymphoma (A- 20) (Edinger *et al*., 2003a). Recipient mice were weighed, given a clinical score, and imaged using the IVIS 50 imaging system three times per week until day 70 or longer. Clinical scores were composed of scores for skin integrity, fur texture, posture, activity, diarrhea, and weight loss. Imaging was done by injecting recipients I.P. with D-luciferin to detect tumor cell bioluminescence. To produce mixed bone marrow chimeras, Thy1.1 mice were lethally irradiated and reconstituted with a 1:4 (WT: *TCF-7 cKO*) mixture of bone marrow cells (total 50x10^6^ cells), then rested for 9 weeks. At 9 weeks, tail vein blood was collected and checked by flow cytometry for CD45.1 and CD45.2 to ensure reconstitution with both donor cell types. At 10 weeks, mice were used for phenotyping experiments.

### Flow Cytometry, Sorting, and Phenotyping

Splenocytes (or cells from other organs) were obtained from WT or *TCF-7* cKO mice or recipient allotransplanted mice. Lymphocytes were obtained and lysed with RBC Lysis Buffer (00-4333-57 from eBioscience) to remove red blood cells if needed. Cells were then stained with extracellular markers for 30 min on ice in MACS buffer (1X PBS with EDTA and 4g/L BSA). If intracellular markers were used, the cells were then fixed and permeabilized using the Fix/Perm Concentrate and Fixation Diluent from FOXP3 Transcription Factor Staining Buffer Set (eBioscience cat. No. 00-5523-00). The cells were then run on a BD LSR Fortessa cytometer and data were analyzed using FlowJo software v9 (Treestar). All antibodies were used at a 1:100 dilution. For FACS sorting, the same methods were applied, and cells were run on a BD FACS Aria IIIu with cold-sorting blocks. Cells were sorted into sorting media (10% FBS in RPMI) or Trizol, depending on the experiment.

Depending on the experiment, antibodies used were: anti-CD4 (FITC, PE, BV785, BV21), anti- CD8 (FITC, PE, APC, PerCP, Pacific Blue, PE/Cy7), anti-CD3 (BV605 or APC/Cy7), anti- H2Kb-Pacific Blue, anti-H2Kd-PE/Cy7, anti-CD122 (FITC or APC), anti-CD44 (APC or Pacific Blue), anti-CD62L (APC/Cy7), anti-TNF-α-FITC, anti-IFN-γ-APC, anti-Eomes (AF488 or PE/Cy7), anti-T-bet-BV421, anti-CD45.2-PE/Dazzle594, anti-CD45.1-APC, anti-Ki67 (PE or BV421), anti-PD1-BV785, anti-CTLA4-PE, NKG2D-BV711, Granzyme B-PE/Cy7.

### Histology

Recipient mice were allotransplanted as described, and organs were removed for histology at day 7, and day 21 post-transplant. Spleen, liver, small intestine, and skin (from ear and back) were fixed, sectioned, and stained with H&E at Cornell University (https://www.vet.cornell.edu/animal-health-diagnostic-center/laboratories/anatomic-pathology/services). A pathologist (L.S) analyzed the sections for T cell-induced damage.

### Cytokine Restimulation

Recipient BALB/c mice were allotransplanted with 1.5X10^6^ CD3 donor T cells and euthanized at day 7. Splenocytes were taken and cultured for 6 hours with GolgiPlug (1:1000) and PBS (control) or anti-CD3 (1ug/mL)/anti-CD28 (2ug/mL) (TCR stimulation) at 37 C and 7% CO2. After 6 hours of culture, the cells were stained for CD3, CD4, CD8, H2K^b^, TNF-α, and IFN-γ using the BD Cytokine Staining kit (BD Biosciences, 555028), and run on a flow cytometer.

### LegendPLEX Serum ELISA Assay

Serum from cardiac blood was collected from recipient mice in the cytokine restimulation experiment. Serum was analyzed using the Biolegend LEGENDPlex Assay Mouse Th Cytokine Panel kit (741043). This kit quantifies serum concentrations of: IL-2 (T cell proliferation), IFN-γ and TNF-α (Th1 cells, inflammatory), IL-4, IL-5, and IL-13 (Th2 cells), IL-10 (Treg cells, suppressive), IL-17A/F (Th17 cells), IL-21 (Tfh cells), IL-22 (Th22 cells), IL-6 (acute/chronic inflammation/T cell survival factor), and IL-9 (Th2, Th17, iTreg, Th9 – skin/allergic/intestinal inflammation).

### Western blot

Splenocytes from WT or *TCF-7* cKO donor mice were collected. CD8 T cells were separated using CD8 MACS beads. CD8 T cells were either stimulated with 2.5ug/ml anti- CD3 (Biolegend #100202) and anti-CD28 antibodies (Biolegend #102115) for 10 minutes or left unstimulated. These cells were counted and lysed with RIPA Buffer (89900 from Thermo Fisher) plus protease inhibitors (11697498001 from Millipore Sigma) and phosphatase inhibitors (P5726-1ML and P0044-1ML from Millipore Sigma). The lysates were run on a Western blot and probed for Perforin (Cell Signaling Technology #3693), Granzyme B (Cell Signaling Technology #4275), LCK (Thermo Fisher PA5-34653), ZAP70(Cell Signaling Technology #3165), LAT(Cell Signaling Technology # 45533), ITK (Thermo Fisher PA5-49363), PLCγ1 (Cell Signaling Technology #2822), ERK1-2 (Cell Signaling Technology #9107), JAK 2(Cell Signaling Technology # 3230), JAK3 (Cell Signaling Technology #8863), STAT3 (Cell Signaling Technology #9139), p65-Rela (Cell Signaling Technology #4764), AKT (Cell Signaling Technology #9272),, and β-actin (Cell Signaling Technology #4970). All the western blots repeated at least three times and one representative of each protein and quantification is shown.

### qPCR Analysis

To perform qPCR, BALB/c mice were allotransplanted as described (1X10^6^ CD3 donor T cells). Pre-transplant donor cells and post-transplant (day 7) spleen and liver cells from recipients were FACS-sorted to obtain CD8 donor cells. The cells were sorted into Trizol, RNA was extracted using chloroform (https://www.nationwidechildrens.org/

Document/Get/93327), and eluted using the Qiagen RNEasy Mini kit (74104 from Qiagen). Concentration was checked with a spectrophotometer, then RNA was converted to cDNA with an Invitrogen Super Script IV First Strand synthesis System kit (18091050 from Invitrogen). Final cDNA concentration was checked with a spectrophotometer, then cDNA was mixed with Taqman Fast Advanced Master Mix (4444557 from Invitrogen) at a 10ng/μL cDNA concentration. This master mix was added to premade 96 well TaqMan Array plates with chemokine/chemokine receptor primers (Thermo Fisher, Mouse Chemokines & Receptors Array plate, 4391524). qPCR was performed in a Quant Studio 3 thermocycler, and data were analyzed using the Design and Analysis software v2.4 (provided by Thermo Fisher). Five separate recipient mice were sorted, and cells were combined to make one sample for qPCR testing per condition/organ.

### NKG2D expression and NKG2D mediated cytotoxicity in CD8 T cells

To determine the NKG2D expression in CD8 T cells, we obtained splenocytes from WT and *TCF-7* cKO mice and stimulated T cells with 2.5ug/ml anti-CD3 (Biolegend #100202) and anti-CD28 antibodies (Biolegend #102115) for 24, 48, or 72 hours in culture, or left them unstimulated. GolgiPlug (1:1000) was added to stimulated samples for each time point, and samples were incubated at 37 C and 7% CO2. After 6 hours of culture, the cells were stained with LIVE/DEAD Aqua and for CD3, CD8, NKG2D, and Granzyme B using the BD Cytokine Staining kit (BD Biosciences, 555028), and run on a flow cytometer. To assess the NKG2D mediated cytotoxicity, we used luciferase-expressing A20 cells as target cells as described earlier. Effector cells (MACS-sorted CD8 T cells from *TCF-7* cKO or WT mice) were incubated in 2.5μg/ml anti-CD3 and anti-CD28 coated plates for 48 hours to induce optimal NKG2D expression. Then effector cells were added at 40:1 effector-to-target ratios and incubated at 37°C for 4 hours with the A20 cells. Anti- NKG2D antibody (10 μg/mL, Bio X Cell #BE0334) or rat IgG1 isotype control antibody (10 μg/mL, Bio X Cell #BE0334) was added and incubated for 30 minutes before washing and plating. Triplicate wells were averaged and percent lysis was calculated from the data using the following equation: % specific lysis = 100 × (spontaneous death bioluminescence – test bioluminescence)/(spontaneous death bioluminescence – maximal killing bioluminescence).

### Exhaustion/Activation Assay

To determine the *in vitro* exhaustion and activation of CD8 T cells, we obtained splenocytes from WT and *TCF-7* cKO mice and either activated them with 2.5ug/ml anti-CD3 (Biolegend #100202) and anti-CD28 antibodies (Biolegend #102115) for 24, 48, or 72 hours in culture, or left them unstimulated, and stained for CD3, CD8, Ki-67, Tox, and PD-1 markers. To assess exhaustion and activation of *in vivo* donor CD8 T cells, recipient mice were allotransplanted as before (1X10^6^ CD3 donor T cells) and euthanized at day 7. Lymphocytes were obtained from spleen, and stained for CD3, CD4, CD8, H2K^b^, TOX, Ki-67 and PD-1 markers.

### DNA Extraction and PCR

Donor mice were genotyped using PCR on DNA extracted from ear punches. At 4 weeks of age mice were ear punched, and DNA was extracted using the Accustart II Mouse Genotyping kit (95135-500 from Quanta Biosciences). Standard PCR reaction conditions and primer sequences from Jackson Laboratories were used for CD4cre. For Tcf7, primer sequences and reaction conditions were obtained from Dr. Jyoti Misra Sen of NIH.

### RNA Sequencing

Recipient mice were allotransplanted as before (1X10^6^ CD3 donor T cells), except that donor CD8 T cells were also FACS-sorted prior to transplant. A sample of sorted donor cells was also saved for pre-transplanted RNA sequencing in Trizol. At day 7 post-transplant, donor CD8 T cells were FACS-sorted back from recipient spleen of *TCF-7* cKO and WT transplanted mice. The cells were all sorted into Trizol, then RNA was extracted and prepped by the Molecular Analysis Core (SUNY Upstate, https://www.upstate.edu/research/facilities/molecular-analysis.php). Paired end sequencing was done with an Illumina NovaSeq 6000 system at the University at Buffalo Genomics Core (http://ubnextgencore.buffalo.edu). For data analysis we used the statistical computing environment R (v4.0.4), the Bioconductor suite of packages for R, and R studio (v1.4.1106). We calculated the transcript abundance by performing pseudoalignment using the Kallisto (Bray et al., 2016) (version 0.46.2). Calculated Transcript per million (TPM) values were normalized and fitted to a linear model by empirical Bayes method with the Voom (Law et al., 2014) and Limma (Ritchie et al., 2015) R packages to determine Differentially expressed genes – DEGs (FDR<0.1, after controlling for multiple testing using the Benjamini-Hochberg method). DEG’s were used for hierarchical clustering and heatmap generation in R. Gene Ontology enrichment analysis was conducted using either the Function Annotation Chart tools using only GO BP and KEGG terms in the Database for Visualization and Integrative Discovery (Huang da et al., 2009a; b) (Huang da *et al*., 2009a) DAVID enrichment scores >1.3 are equivalent to a P value<0.05. For Gene set enrichment analysis (GSEA) we used Hallmark and C2 gene set collections of Molecular Signatures Database (MsigDB) and cluster Profiler package in R. Data will be deposited on the Gene Expression Omnibus (GEO) database for public access https://www.ncbi.nlm.nih.gov/geo/

The RNAseq experiment described here was performed as part of the experiment described in other recent publications from our laboratory (Mammadli et al., 2021a; Mammadli et al., 2021b; Mammadli et al., 2020; Mammadli et al., 2021c). Therefore, the data generated for WT pre- and post-transplanted samples (CD4 and CD8) are the same as that shown in the papers mentioned, but here, these data are compared to data for *Cat-Tg* mice(Mammadli *et al*., 2021a).

### Statistical Analysis

Unless otherwise noted in the figure legends, all numerical data are presented as means and standard deviations with or without individual points. Analysis was done in GraphPad Prism v7 or v9. Most data were analyzed with Student’s t-test, one-way ANOVA, or two-way ANOVA, with Tukey’s multiple comparisons test for ANOVA methods, depending on the number of groups. Kaplan-Meier survival analyses were done for survival experiments. All tests were two-sided, and p-values less than or equal to (≤) 0.05 were considered significant. Transplant experiments used 3-5 mice per group, with at least two repeats. *Ex vivo* experiments were done two to three times unless otherwise noted with at least three replicates per condition each time. RNA seq was done once with three replicates per group and condition. qPCR was done once with one sample per condition, and 5 mice were combined to make the one sample. Western blots were done 3 times for unstimulated and 10 min anti-CD3/CD28 stimulated samples, one experiment each is shown.

### Study Approval

All animal studies were reviewed and approved by the IACUC at SUNY Upstate Medical University. All procedures and experiments were performed according to these approved protocols.

## Author Contributions

RH, MM, and MK designed and conducted experiments, analyzed data, and wrote the manuscript. MK assisted with scientific/technical research design. MK, and JMS edited the manuscript. MK, SH assisted with data collection. LS performed histology analyses. QY provided technical and scientific advice and assisted with data analysis.

## Acknowledgments

We thank all members of the Karimi lab for helpful discussions. We also thank Joel Wilmore for help in flow cytometry analysis. This research was funded in part by a grant from the National Blood Foundation Scholar Award to MK, the National Institutes of Health (NIH LRP #L6 MD0010106 and K22 (AI130182) to MK), and an Upstate Medical University Cancer Center grant (1146249-1-75632) to MK.

JMS was supported by the Intramural Research Program of the National Institute of Aging. We thank Dr. Howard Xue for permission to use *TCF-7* cKO mice. *TCF-7* flox/flox mice were provided by Dr. Jyoti Misra Sen from NIH. RH was a PH. D student at SUNY Upstate Medical University at the time the study was conducted from 2017-2021. A version of this manuscript was previously included as a chapter in RH’s dissertation.

---

**Summary Figure.**
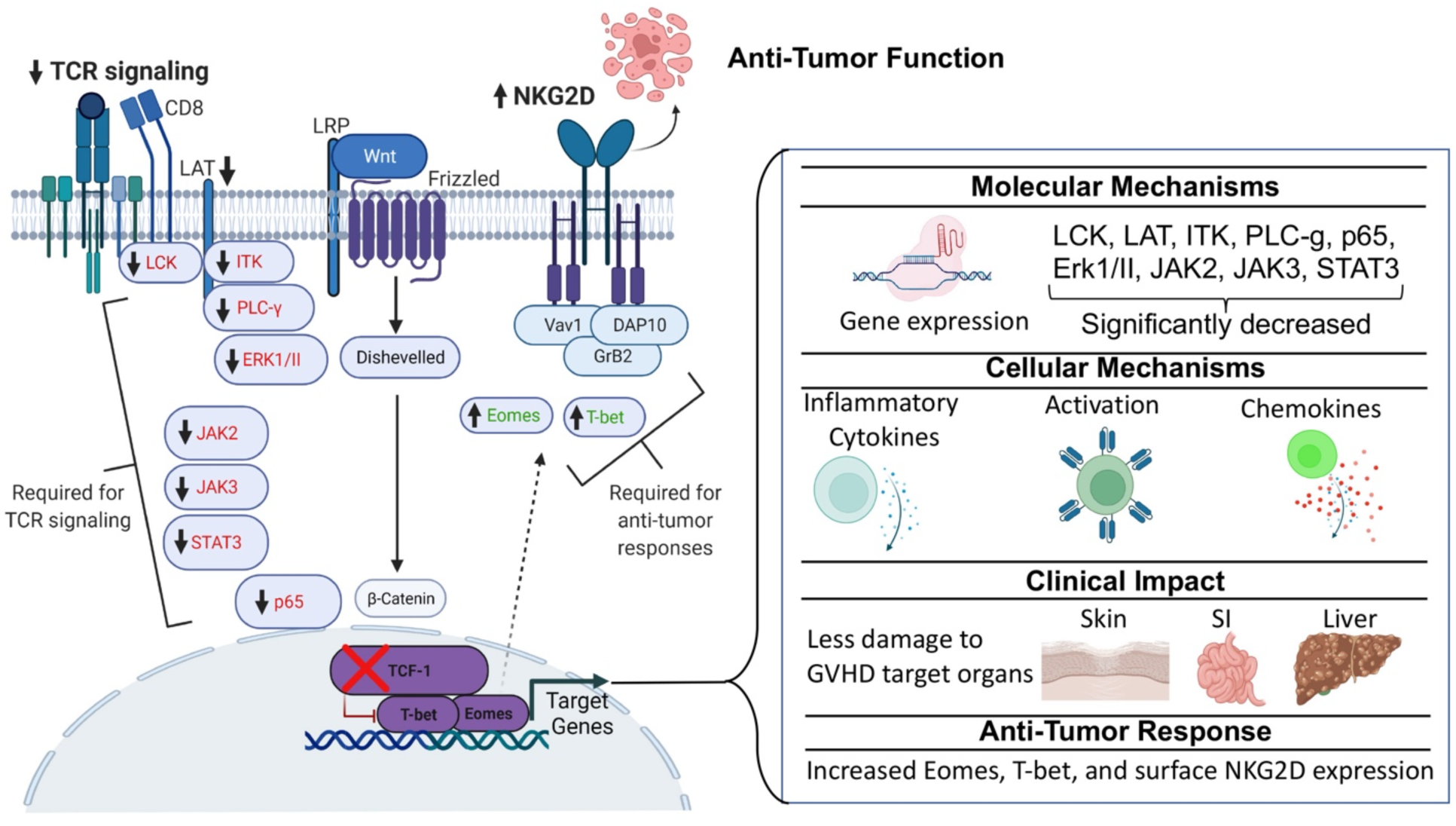
*TCF-7* is required for persistent function of CD8 T cells but dispensable for anti-tumor response. Here we have utilized a novel mouse model that lacks *TCF-7* specifically on CD8 T cells for an allogeneic transplant model. We uncovered a molecular mechanism of how *TCF-7* regulates key signaling pathways at both transcriptomic and protein levels. These key molecules included LCK, LAT, ITK, PLC-γ1, p65, ERK I/II, and JAK/STAT signaling. Next, we showed that the lack of *TCF-7* impacted phenotype, proinflammatory cytokine production, chemokine expression, T cell activation. We provided clinical evidence for how these changes impact GVHD target organs (skin, small intestine, and liver). Finally, we provided evidence that *TCF-7* regulates NKG2D expression on mouse naïve and activated CD8 T cells. We have shown that CD8 T cells from *TCF-7* cKO mice mediate cytolytic functions via NKG2D.

**Supp.Fig.1.**
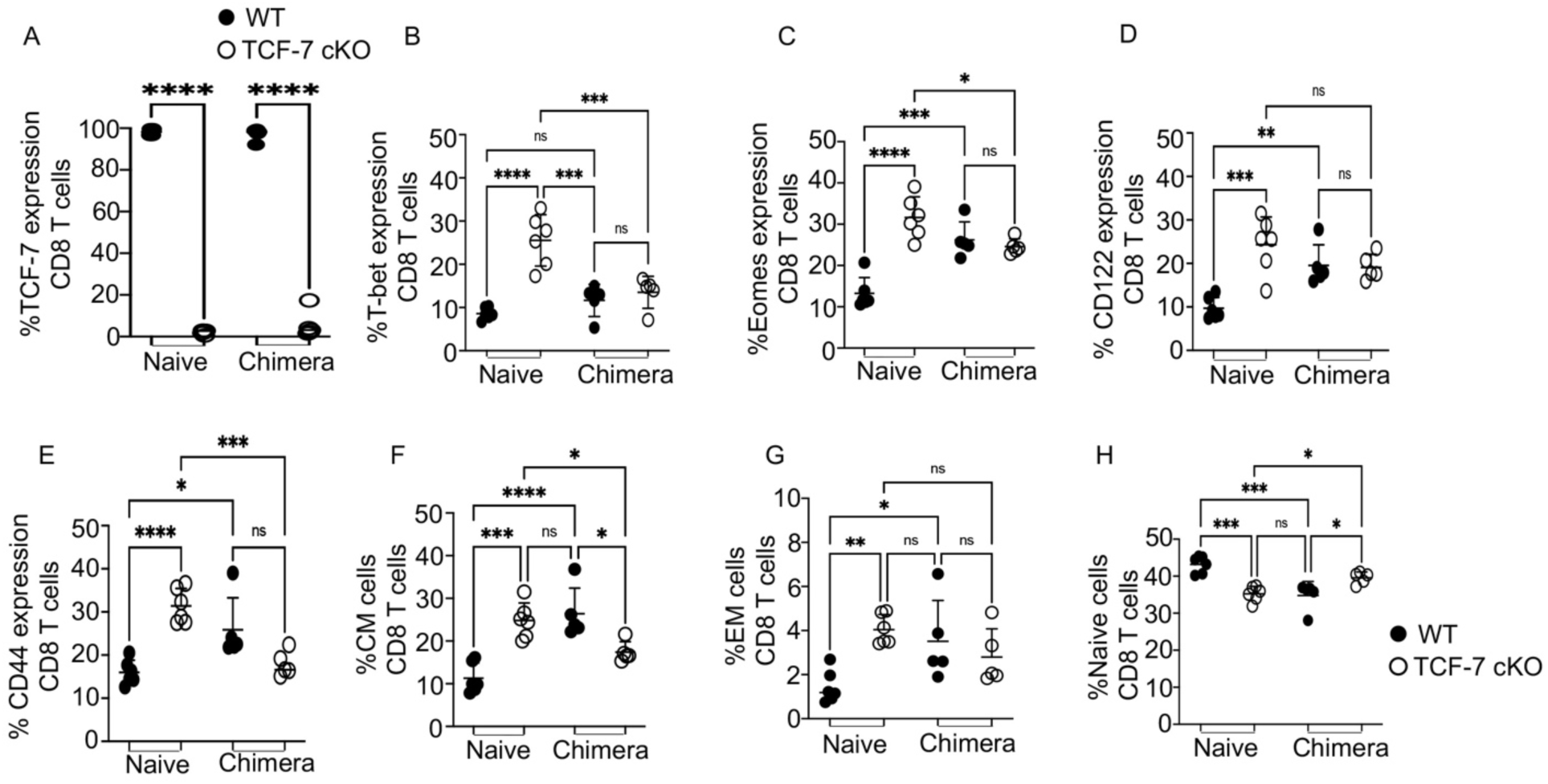
Related to Fig2. Loss of *TCF-7* drives changes to mature CD8 T cell phenotype that are cell-intrinsic, with the possibility of extrinsic effects. Bone marrow chimeras were developed by lethally irradiating Thy1.1 mice and reconstituting with a 1:4 (WT:*TCF-7* cKO) mixture of bone marrow cells. Blood was tested at 9 weeks to ensure reconstitution with both donor cell types, and splenocytes were used at 10 weeks for phenotyping by flow cytometry. (A) Percentage of CD8 T cells from chimeric and naive mice expressing *TCF-7*. (B) Percentage of CD8 T cells from chimeric and naive mice expressing T-bet. (C) Percentage of CD8 T cells from chimeric and naive mice expressing Eomes. (D) Percentage of CD8 T cells from chimeric and naive mice expressing CD122. (E) Percentage of CD8 T cells from chimeric and naive mice expressing CD44. (F) Percentage of CD8 T cells from chimeric and naive mice expressing central memory (CM) phenotype. (G) Percentage of CD8 T cells from chimeric and naive mice expressing effector memory (EM) phenotype. (H) Percentage of CD8 T cells from chimeric and naive mice expressing naïve phenotype. All data are plotted as individual points with mean and SD, all were analyzed with one-way ANOVA, or Student’s t-test (depending on groups). For all graphs, * means p-value ≤ 0.05, *** means p-value ≤ 0.001, and **** means p-value ≤ 0.0001. For naïve cells 3 different experiments combined (N=2-3 per group of mice) and for chimera cell (N=5) with one experiment shown (done once).

**Supp.Fig.2.**
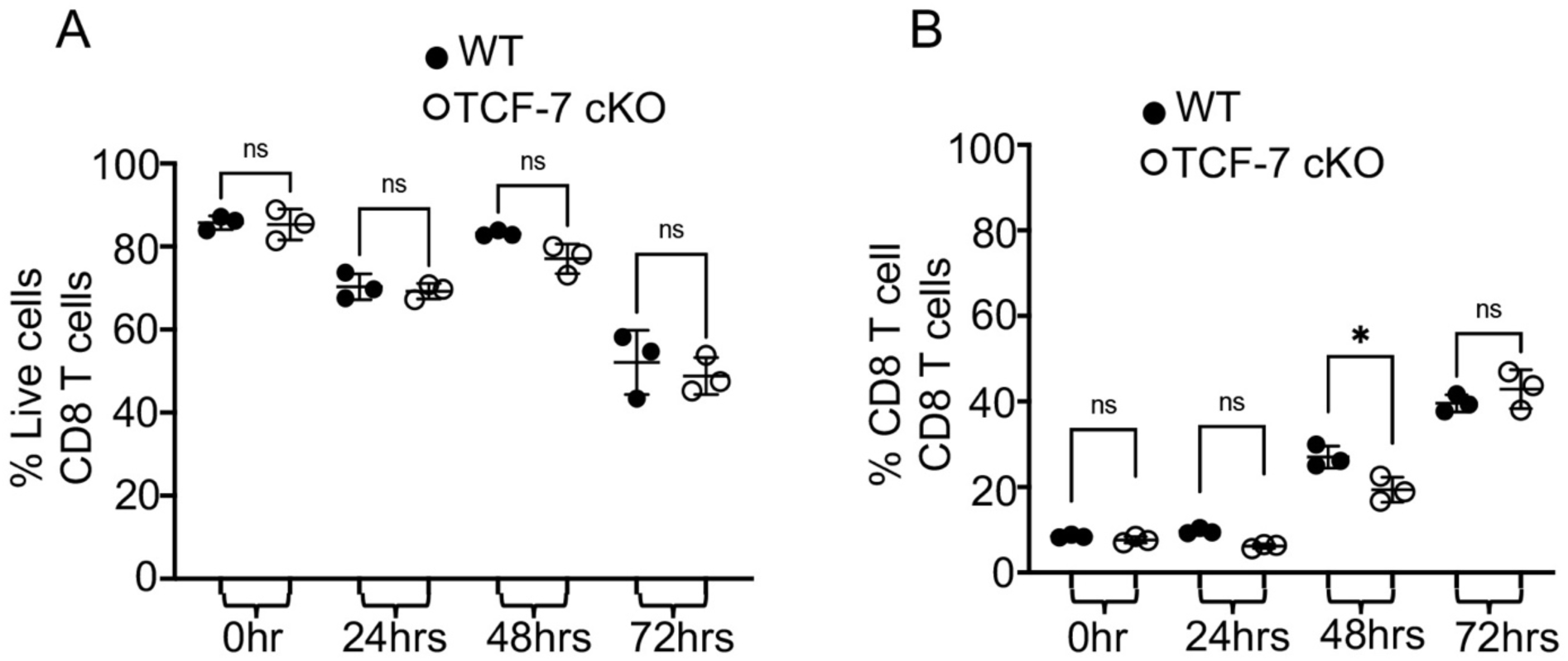
Related to Fig.3. Cell viability and CD8 + T cell percent in NKG2D induction *in vitro*. Total splenocytes were isolated from *TCF-7* cKO or WT mice and either left unstimulated or stimulated with anti-CD3/CD28 for 24, 48, or 72 hrs in culture. GolgiPlug (1:1000) was added to stimulated samples for each time point except 0 hr samples, and samples were incubated at 37 C with 7% CO2. After 6 hours of culture, the cells were stained for CD3, CD8, NKG2D, and Granzyme B expression, as determined by flow cytometry**. (A)** Quantification of percent of live cells (dead cells that were positive for LIVE/DEAD Aqua excluded) for different time points of stimulation. **(B)** Quantification of CD8 T cell percentages for different time points of stimulation. N=4 per group with one representative of 2 experiments shown. All data are shown as individual points with mean and SD, and were analyzed with two-way ANOVA (depending on groups). * means p-value ≤ 0.05, ** means p-value ≤ 0.01, and *** means p-value ≤ 0.001.

**Supp.Fig.3.**
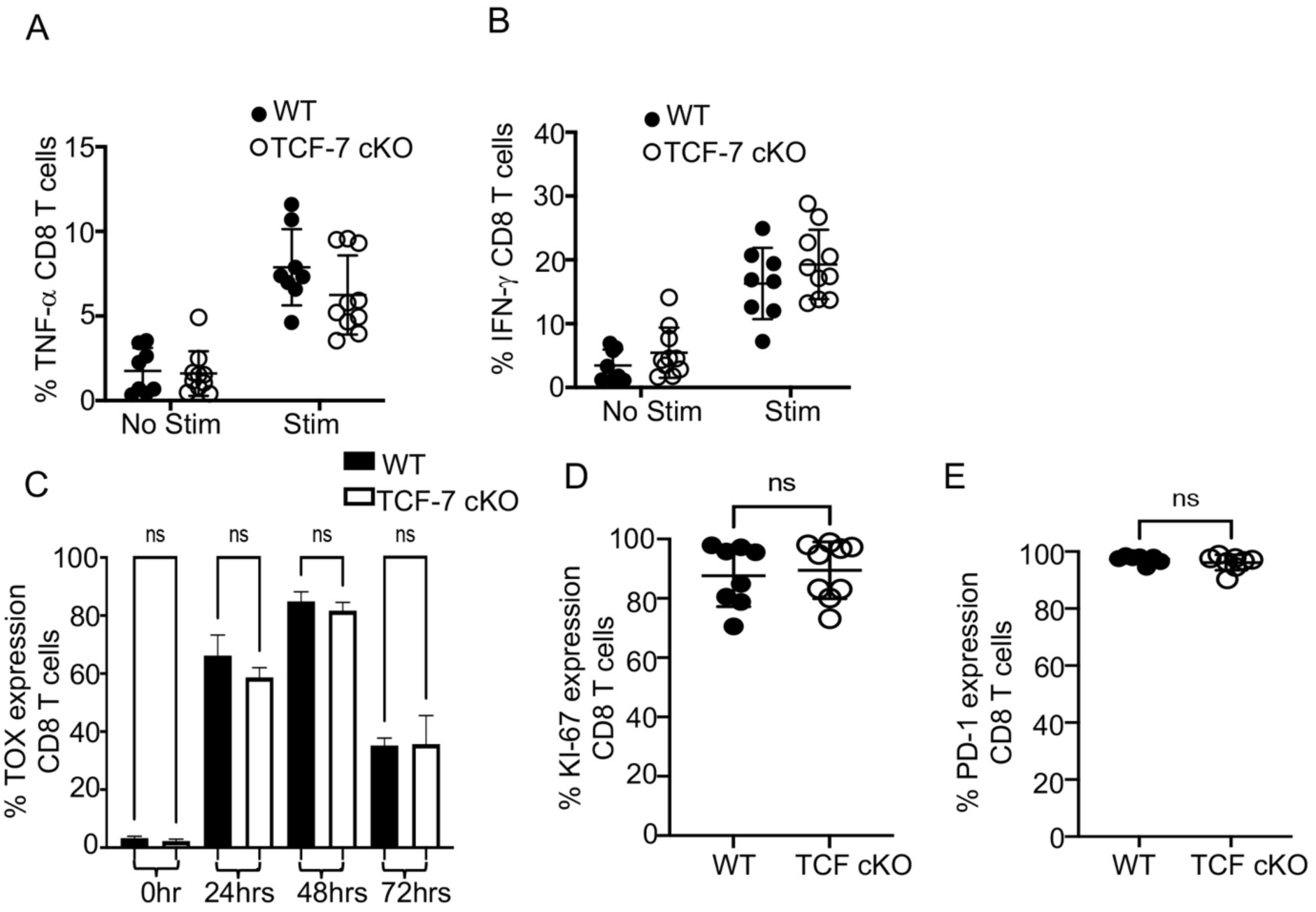
Related to Fig.5. *TCF-7* controls cytokine production and exhaustion of mature alloactivated CD8 T cells. **(A-B)** Recipient mice were allotransplanted with 1.5X10^6^ WT or *TCF-7* cKO donor CD3 T cells as before. Splenocytes were taken at day 7 post-transplant, restimulated by 6-hour culture with Golgi Plug and PBS (control) or anti-CD3/anti-CD28 (TCR restim.), then stained for H2K^b^, CD3, CD4, CD8, TNF-α, and IFN-γ. **(A)** TNF-α production and (**B**) IFN-γ production by donor CD8 T cells, as measured by percent cytokine-positive donor cells. **(C)** Splenocytes from *TCF-7* cKO and WT mice were obtained and either stimulated with anti-CD3/CD28 antibodies for 24hrs, 48hrs, or 72hrs in culture and stained for TOX, or were immediately stained after isolation without stimulation. Percent expression of TOX in CD8 T cells after 72 hours of CD3/CD28 stimulation in culture, as determined by flow cytometry. **(D-E)** Balb/c mice were allo-transplanted as before, with WT BM and WT or *TCF-7* cKO CD8 donor T cells. On day 7, spleens were removed from recipients, processed to isolate lymphocytes, and lymphocytes were stained for CD3, CD8, H2K^b^, Ki-67, and PD-1 to identify proliferating and exhausted T cells, respectively. **(D)** Quantification of the Ki-67 expression in *in vivo* donor CD8 T cells. (**E)** Quantification of the PD-1 expression in *in vivo* donor CD8 T cells. N=3-5 per group for **A-B and D-E** with two experiments shown. N=4 per group for **C,** one representative of two experiments shown. All data are shown as individual points with mean and SD, and were analyzed with Student’s t-test or two-way ANOVA (depending on groups). * means p-value ≤ 0.05, ** means p-value ≤ 0.01, and *** means p-value ≤ 0.001.

**Supp.Fig.4.**
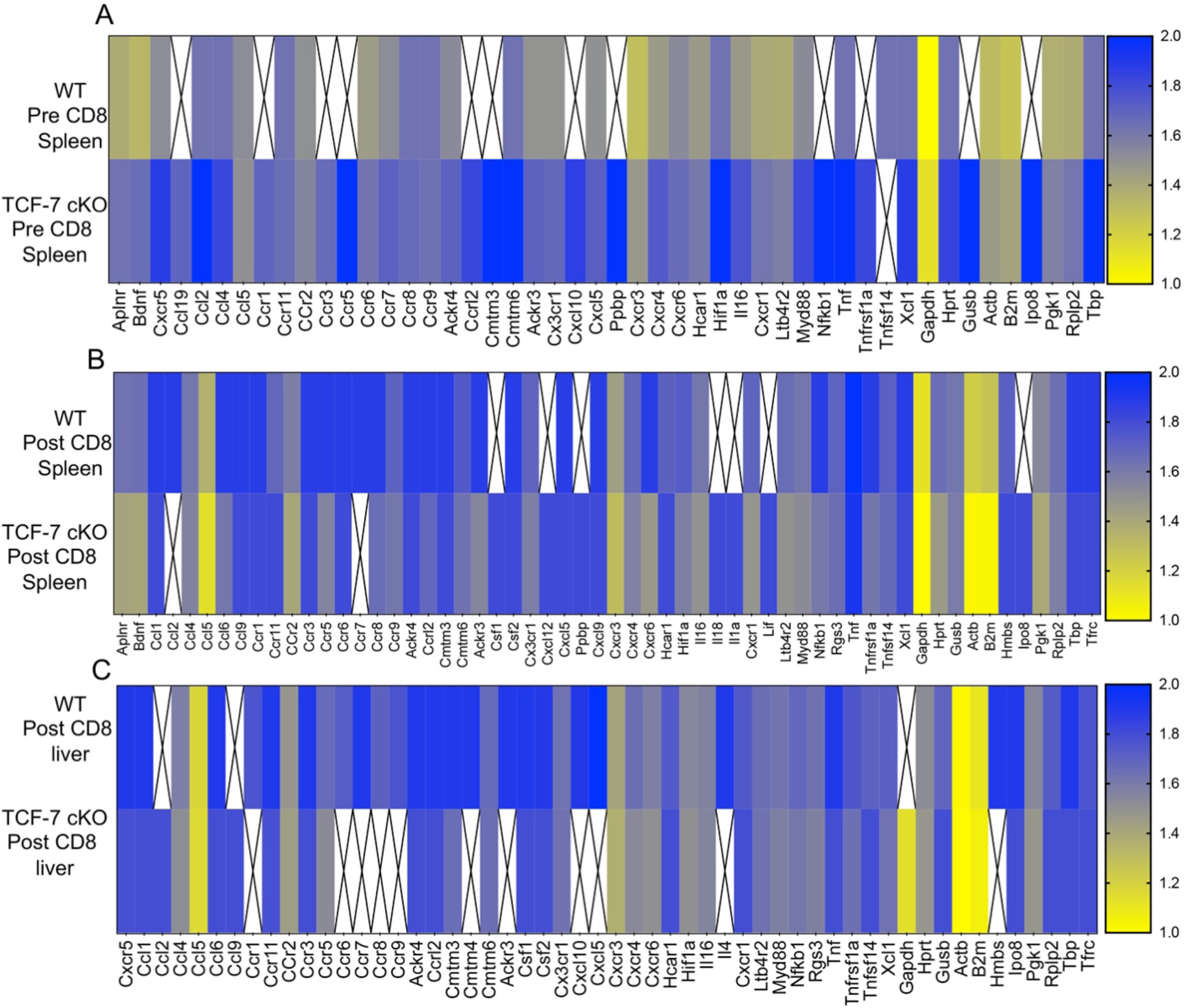
*TCF-7* controls chemokine and chemokine receptor expression of donor CD8 T cells during alloactivation. As before, BALB/c mice were allotransplanted with BM and 1X10^6^ donor CD3 T cells from WT or *TCF-7* cKO mice (not mixed). Donor CD8 T cells were FACS-sorted from spleen of donor’s pre-transplant, and from spleen and liver of recipients at day 7 post-transplant. Cells were sorted into Trizol, then RNA was extracted using chloroform and converted to cDNA for qPCR analysis. cDNA was run on premade mouse chemokine/chemokine receptor assay plates, and results are displayed as heatmaps. Scales are shown at right, with fold change per gene compared to an 18S reference gene on each plate. White boxes with an “X” represent signals too low to detect or otherwise unreadable due to technical limitation/error. **(A)** Pre-transplant spleen donor CD8 T cells, **(B)** post-transplant spleen donor CD8 T cells, and **(C)** post-transplant liver donor CD8 T cells for WT versus *TCF-7* cKO donors. N=5 mice into one sample per condition, summary data shown.

**Supp.Fig.5:**
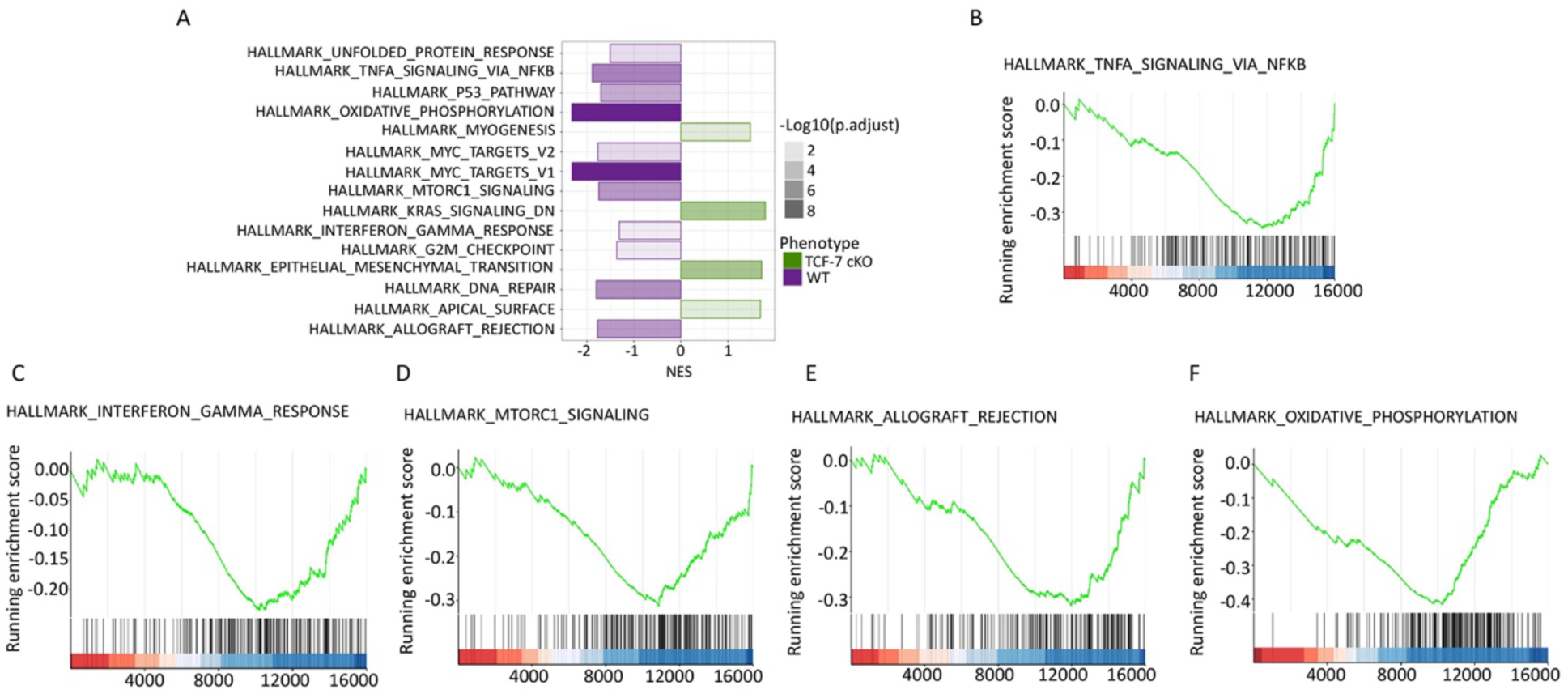
Loss of *TCF-7* alters the enrichment of gene sets in pre-transplanted CD8+T cells. **(A)** Bar plot of the altered pathways indentified in Gene set enrichment analysis (GSEA) by using Hallmark gene set collections of the Molecular Signatures Database (MSigDB) in pre-transplanted CD8 T cells (*TCF-7* cKO compared to WT). The normalized enrichment score (NES) for the pathway is defined as the peak score furthest from zero, with a negative NES meaning enrichment in the WT group. **(B)** GSEA plot for the “HALLMARK_TNFA_SIGNALING_VIA_NFKB” pathway comparing pre-transplanted CD8 T cells from *TCF-7* cKO to WT cells. The running enrichment score (ES) for the pathway is defined as the peak score furthest from zero, with a negative ES meaning enrichment in the WT group. **(C)** GSEA plot for the “HALLMARK_INTERFERON_GAMMA_RESPONSE” pathway comparing pre-transplanted CD8 T cells from *TCF-*7 cKO to WT cells. **(D)** GSEA plot for the “HALLMARK_MTORC1_SIGNALING” pathway comparing pre-transplanted CD8 T cells from *TCF-7* cKO to WT cells. **(E)** GSEA plot for the “HALLMARK_ALLOGRAFT_REJECTION” pathway comparing pre-transplanted CD8 T cells from *TCF-7* cKO to WT cells. **(F)** GSEA plot for the “HALLMARK_OXIDATIVE_PHOSPHORYLATION” pathway comparing pre-transplanted CD8 T cells from *TCF-7* cKO to WT cells.

**Supp.Fig.6.**
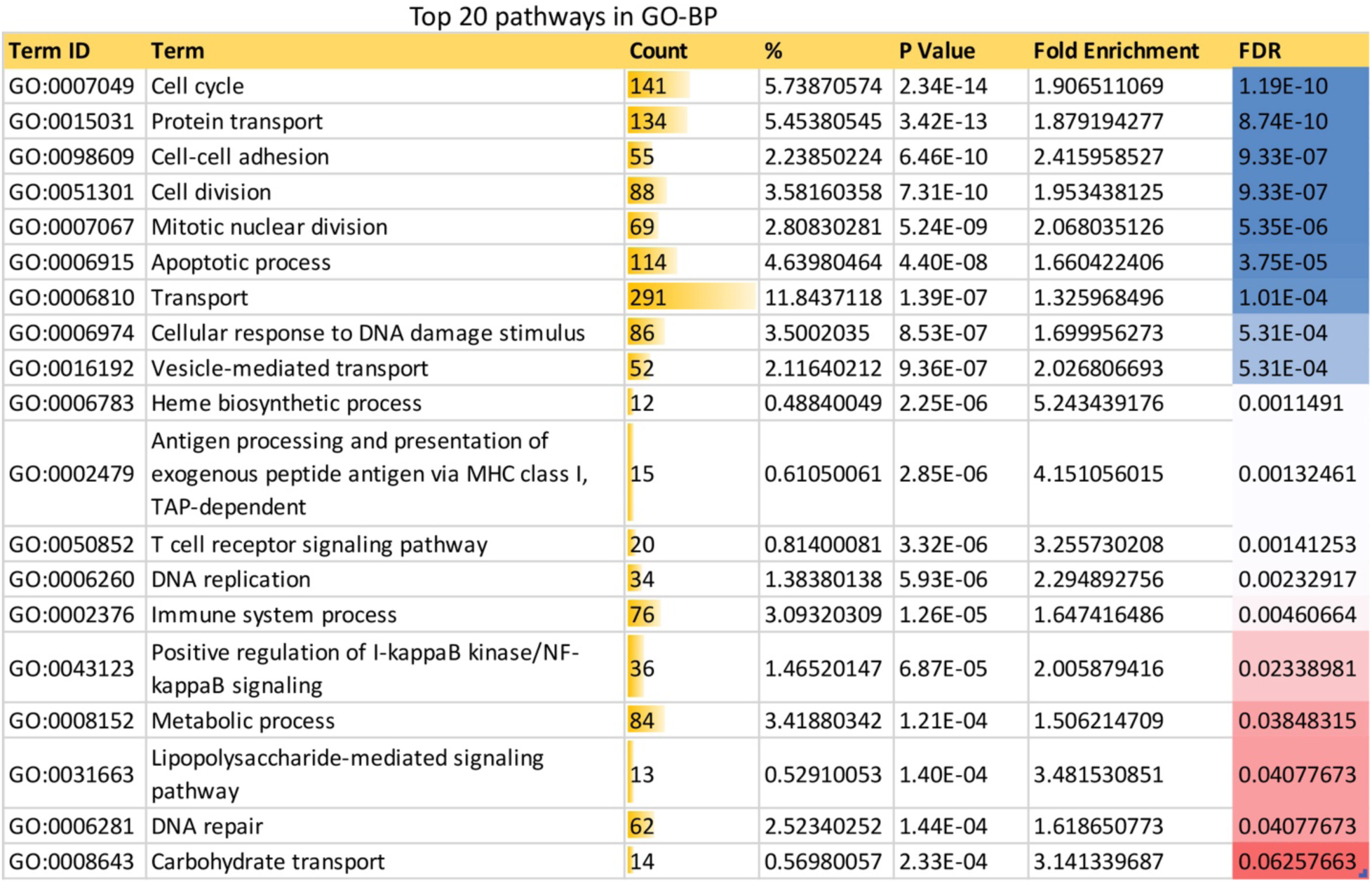
Related to Fig.7. Top 20 GO-BP terms identified in DAVID functional annotation analysis of differentially expressed genes in post-transplanted samples. Count mean – number of genes involved in the Term, % - percentage of involved genes/ total genes in the Term.

**Supp.Fig.7.**
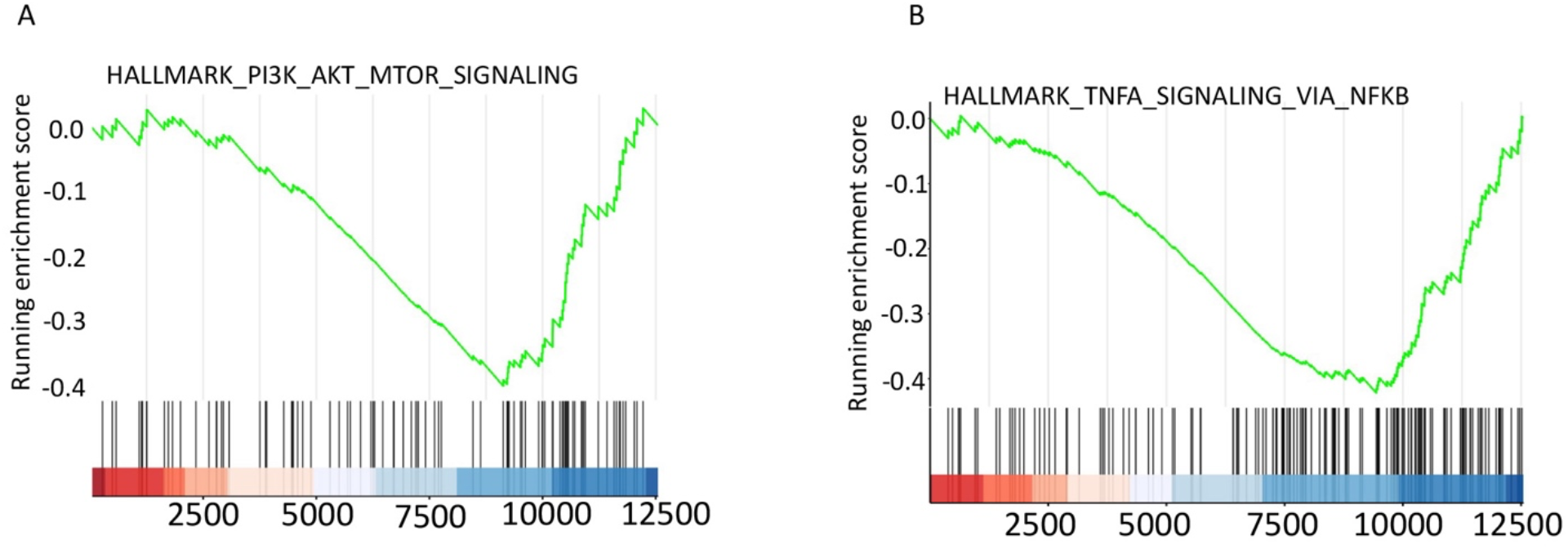
Loss of *TCF-7* alters the Gene Set Enrichment Analysis (GSEA) of post- transplanted CD8+T cells. **(A)** GSEA plot for the “HALLMARK_PI3K_AKT_MTOR_SIGNALING” pathway comparing post-transplanted CD8 T cells from *TCF-7* cKO to WT mice. The running enrichment score (ES) for the pathway is defined as the peak score furthest from zero, with a negative ES meaning enrichment in the WT group. **(B)** GSEA plot for the “HALLMARK_TNFA_SIGNALING_VIA_NFKB” pathway comparing post-transplanted CD8 T cells from *TCF-7* cKO to WT mice. Again, the running enrichment score (ES) for the pathway is defined as the peak score furthest from zero, with a negative ES meaning enrichment in the WT group.

**Supp.Fig.8.**
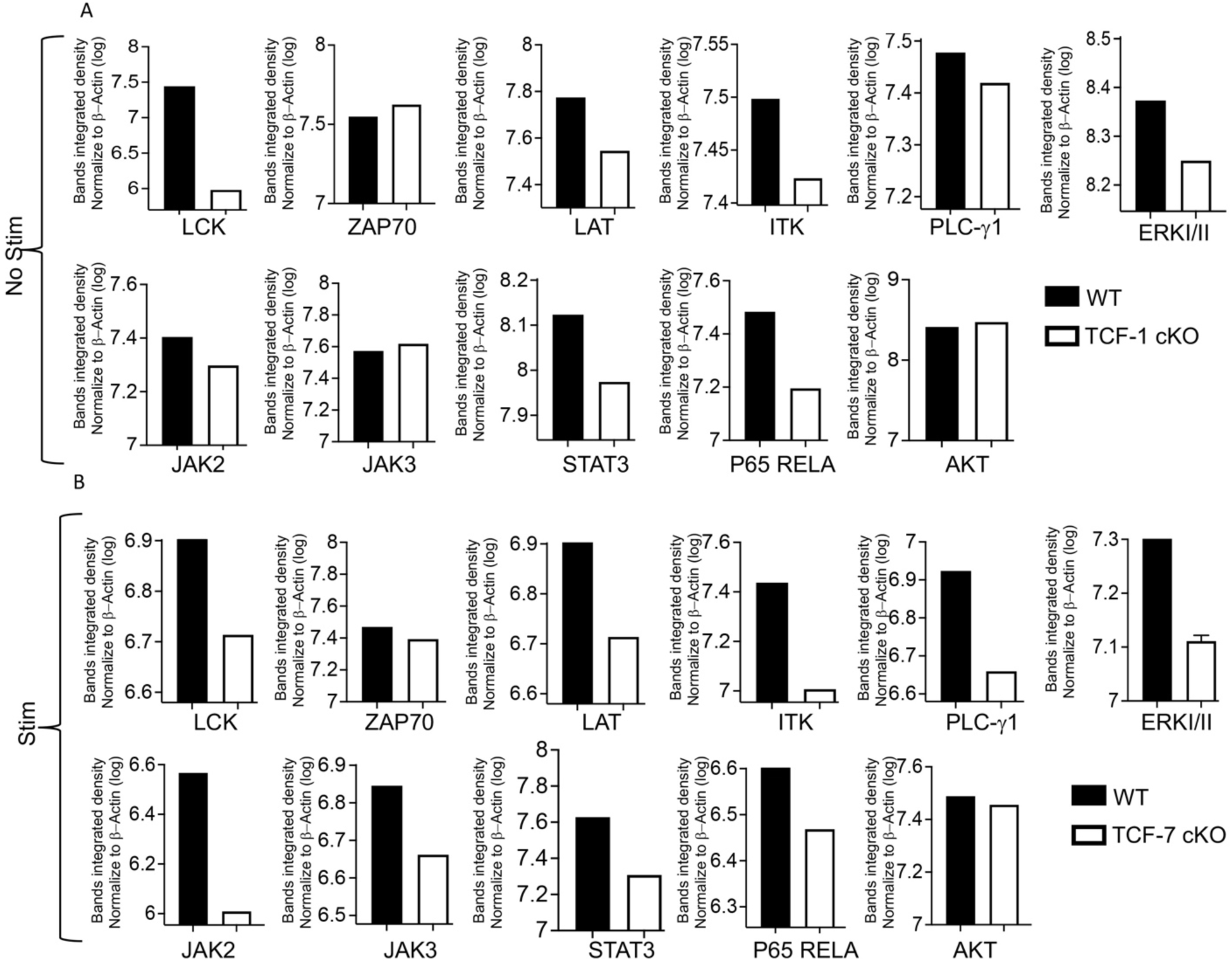
Related to Fig.8. Quantification of Western blot of TCR and JAK-STAT signaling. (**A)** Comparison of the quantified bands integrated density normalized to β-actin for unstimulated CD8 T cells from *TCF-7* cKO and WT mice**. (B)** Comparison of the quantified bands integrated density normalized to β-actin for 10-minute-anti-CD3/CD28-stimulated CD8 T cells from *TCF-7* cKO and WT mice. All the western blots repeated at least three times and one representative of each protein and quantification is shown.

