## Supplementary figures and images for "*TCF-7* is dispensable for anti-tumor response but required for persistent function of mature CD8 T cells"

### Sup.Fig2

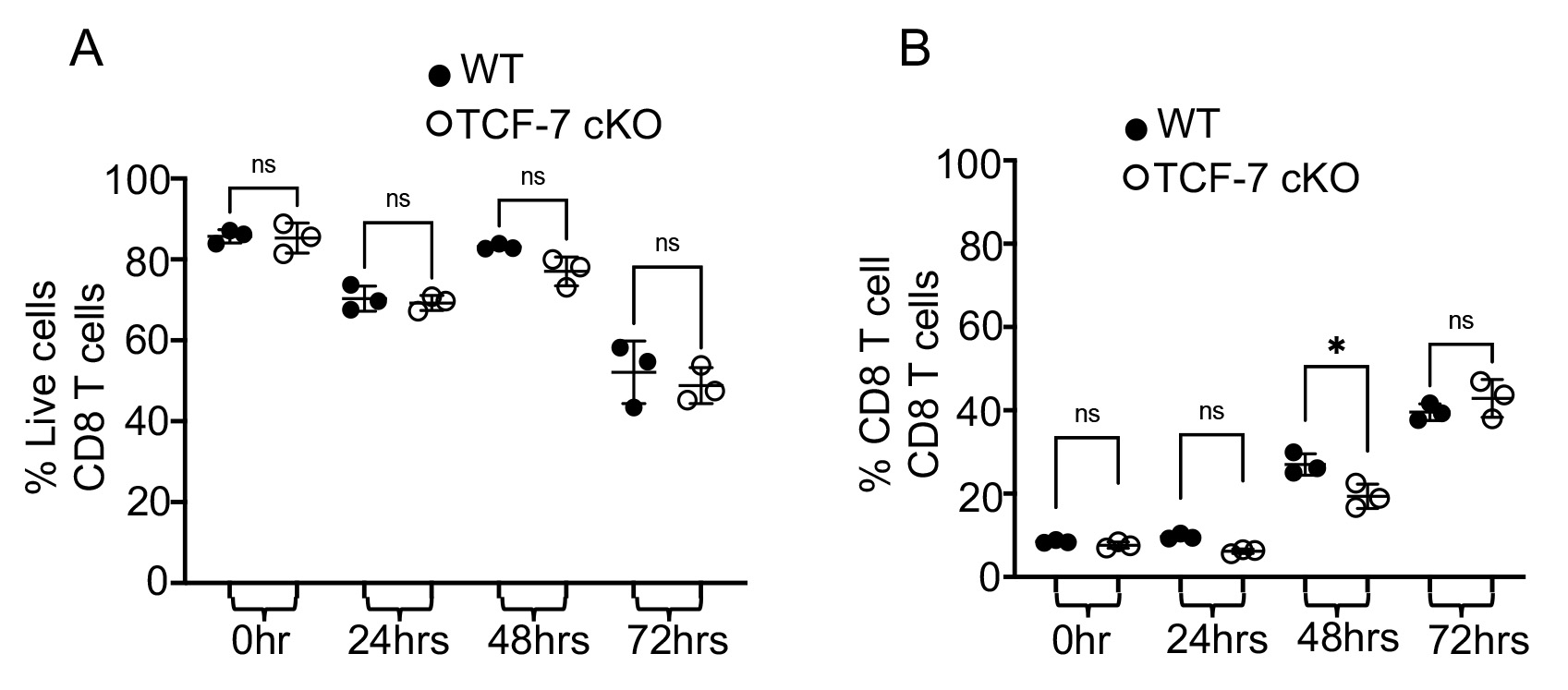

### Sup.Fig.4

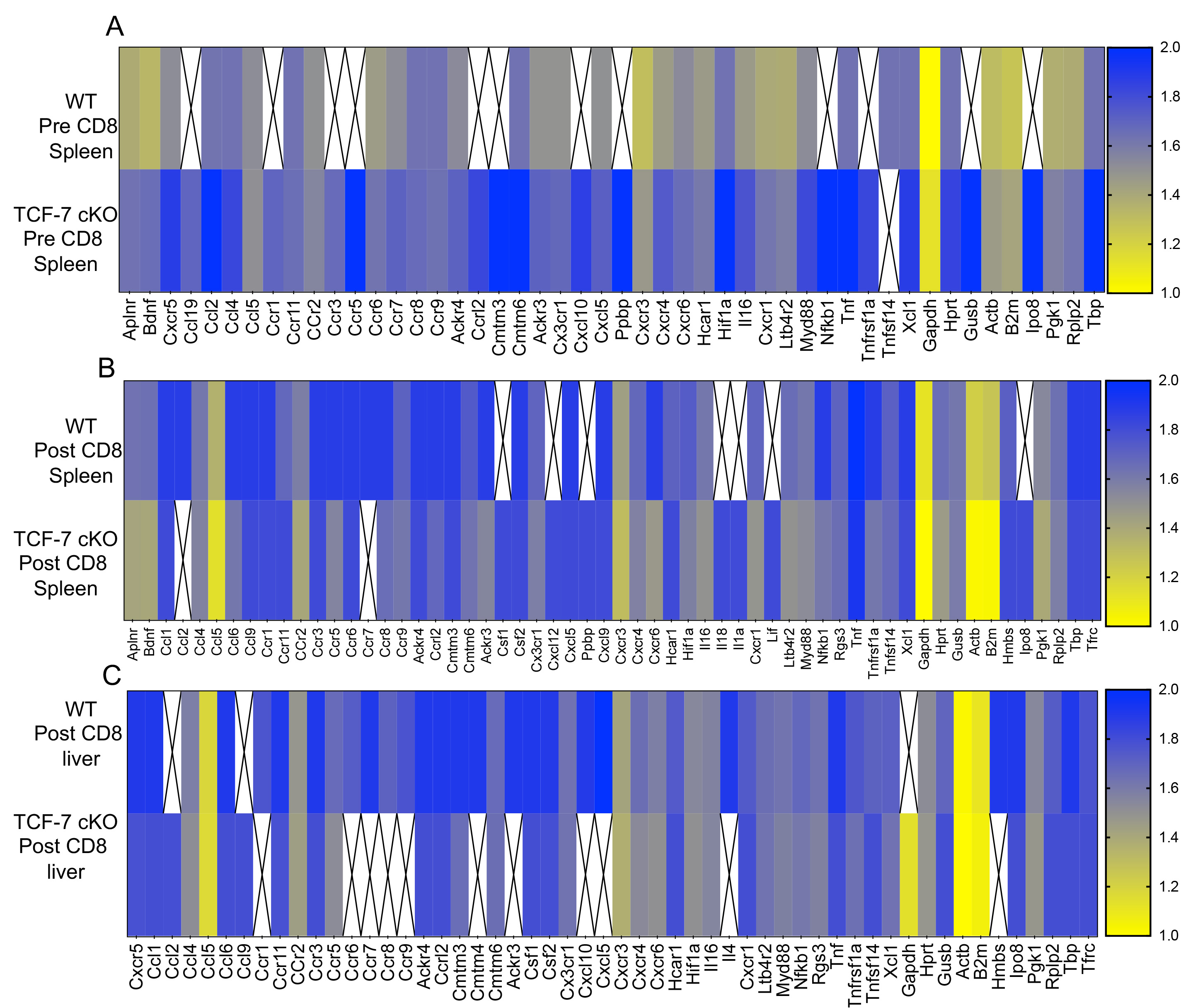

### Sup.Fig.5

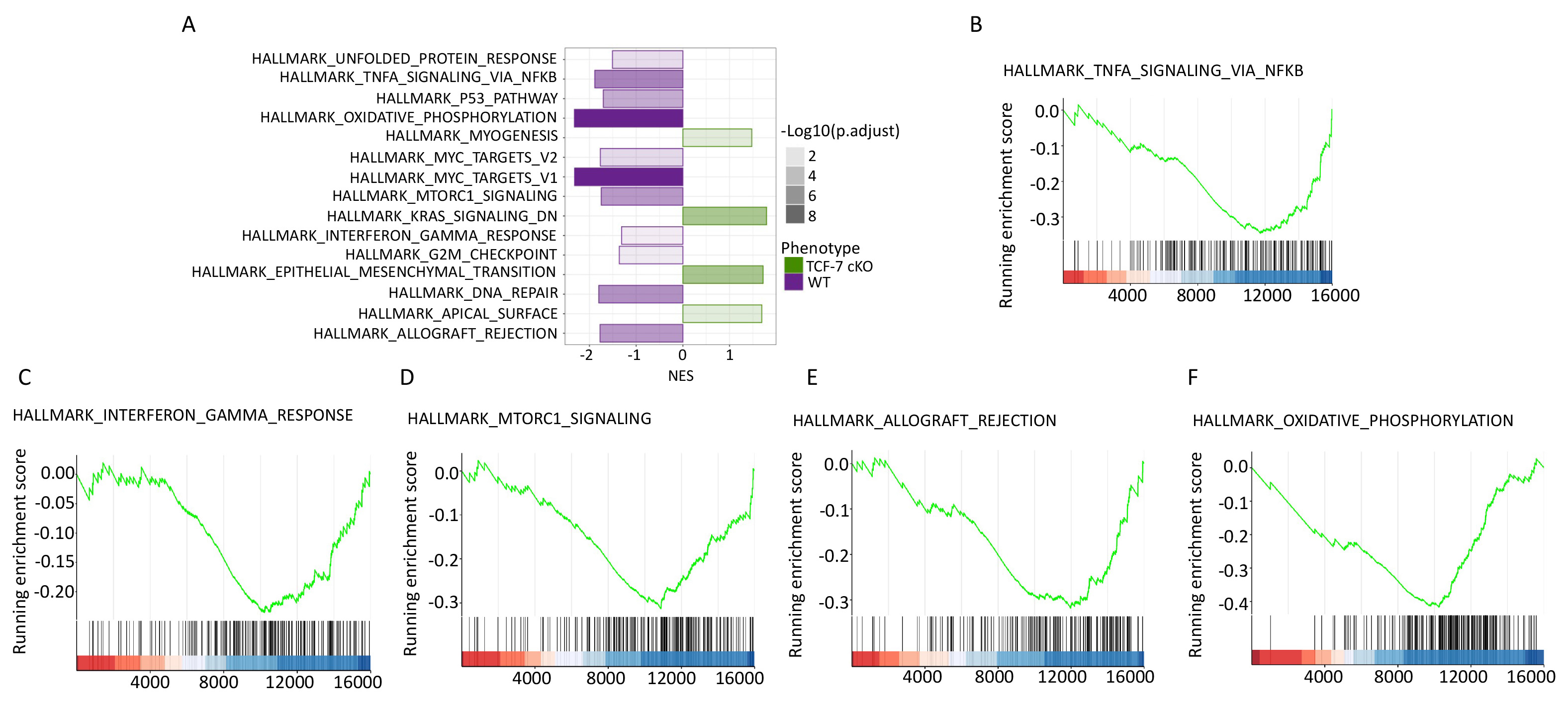

### Sup.Fig.6

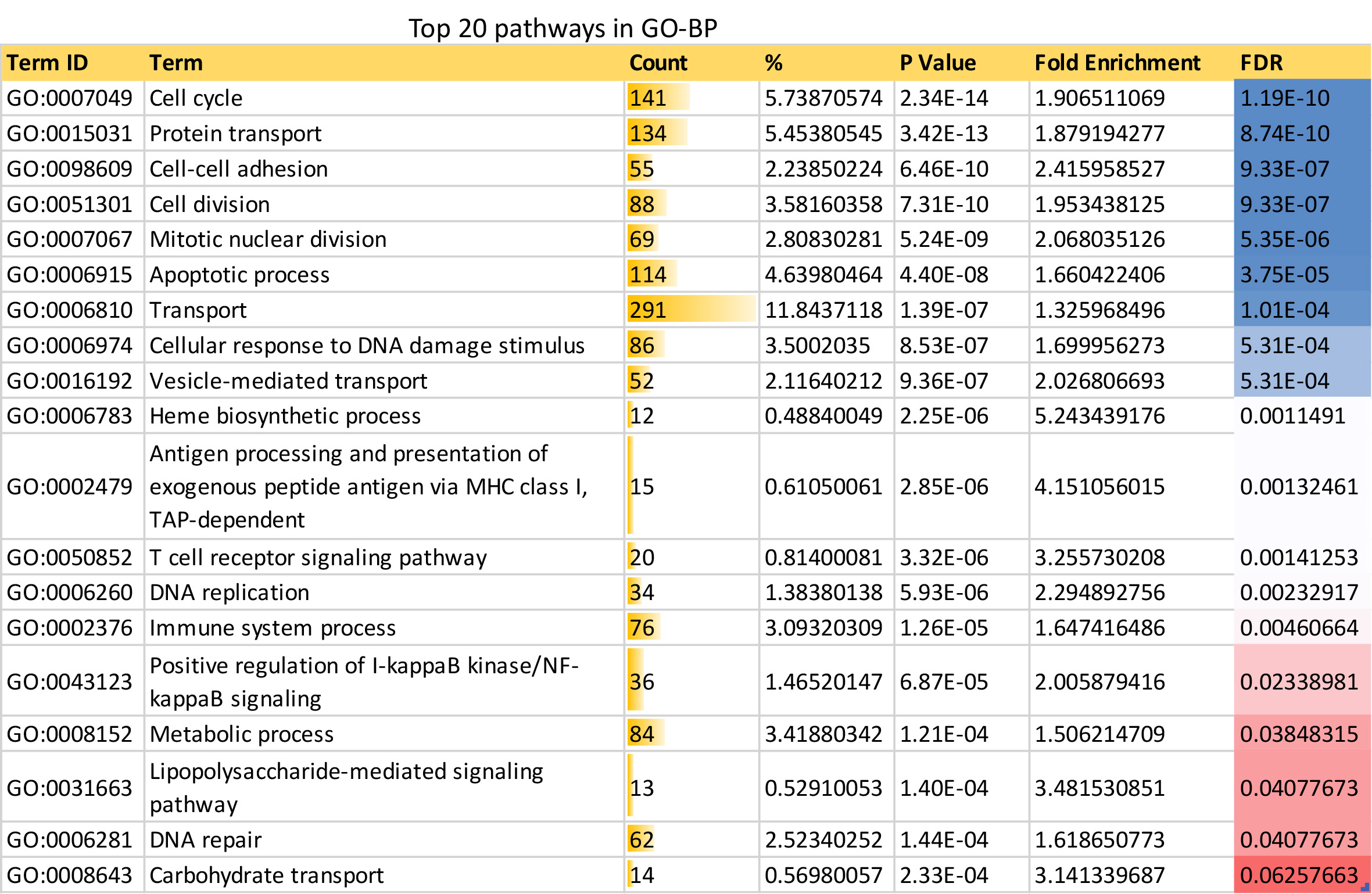

### Sup.Fig.7

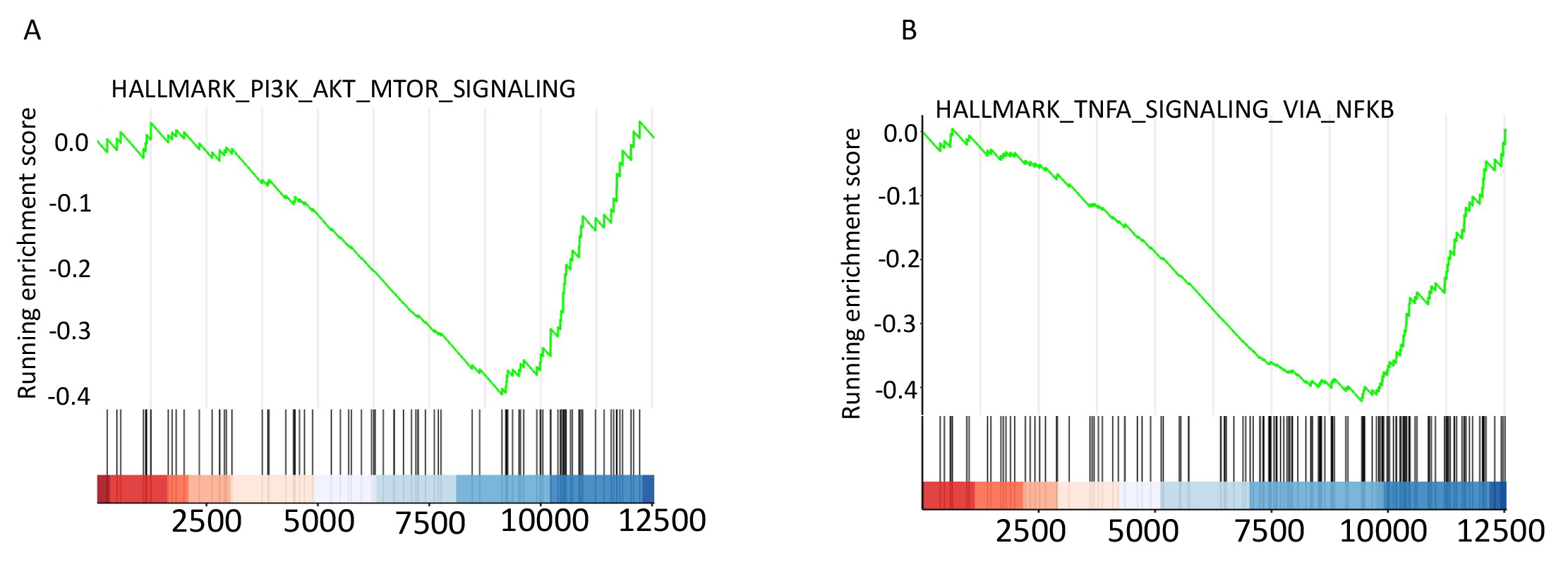

### Sup.Fig.8

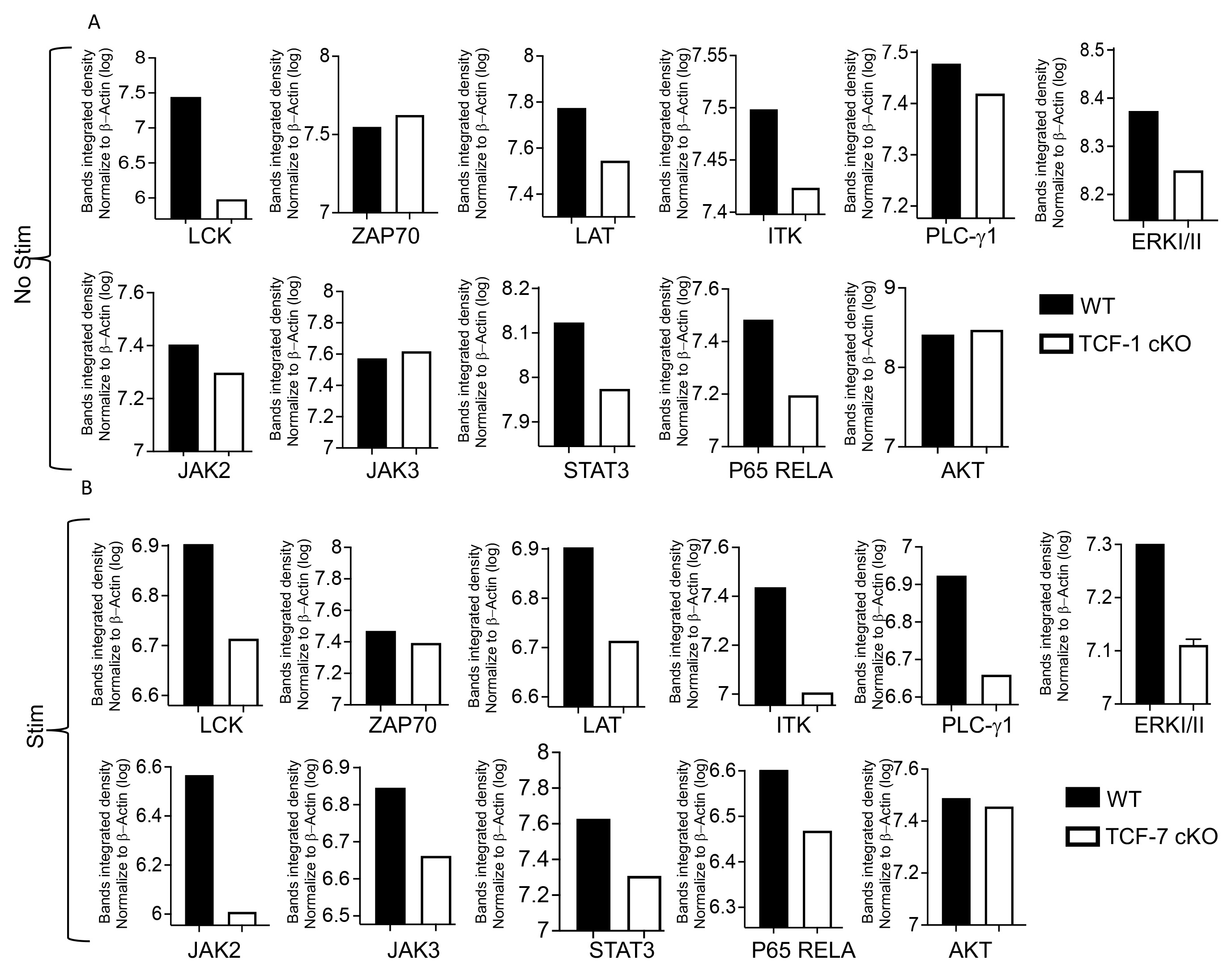

### Sup.Figure1

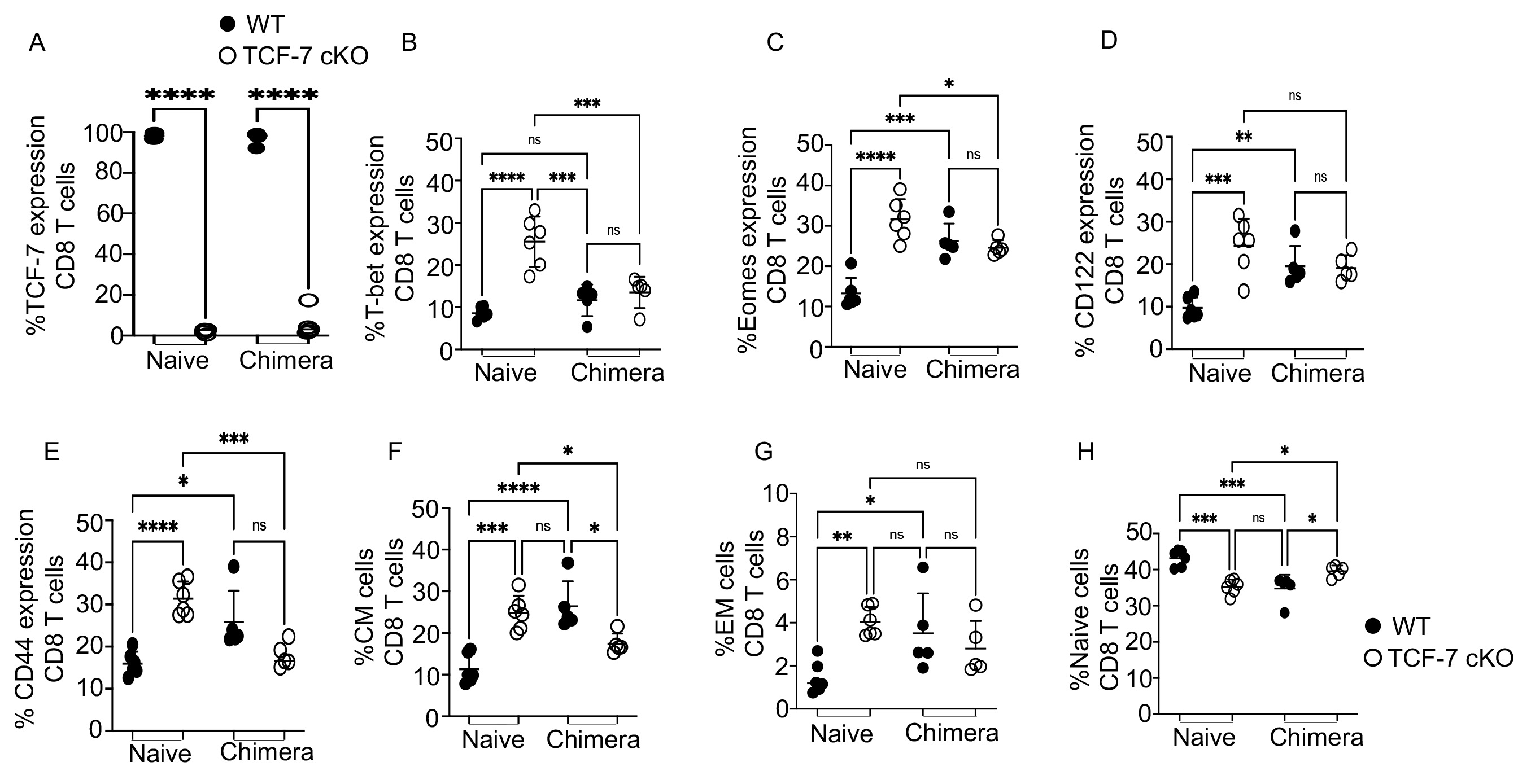

### Sup.Figure3

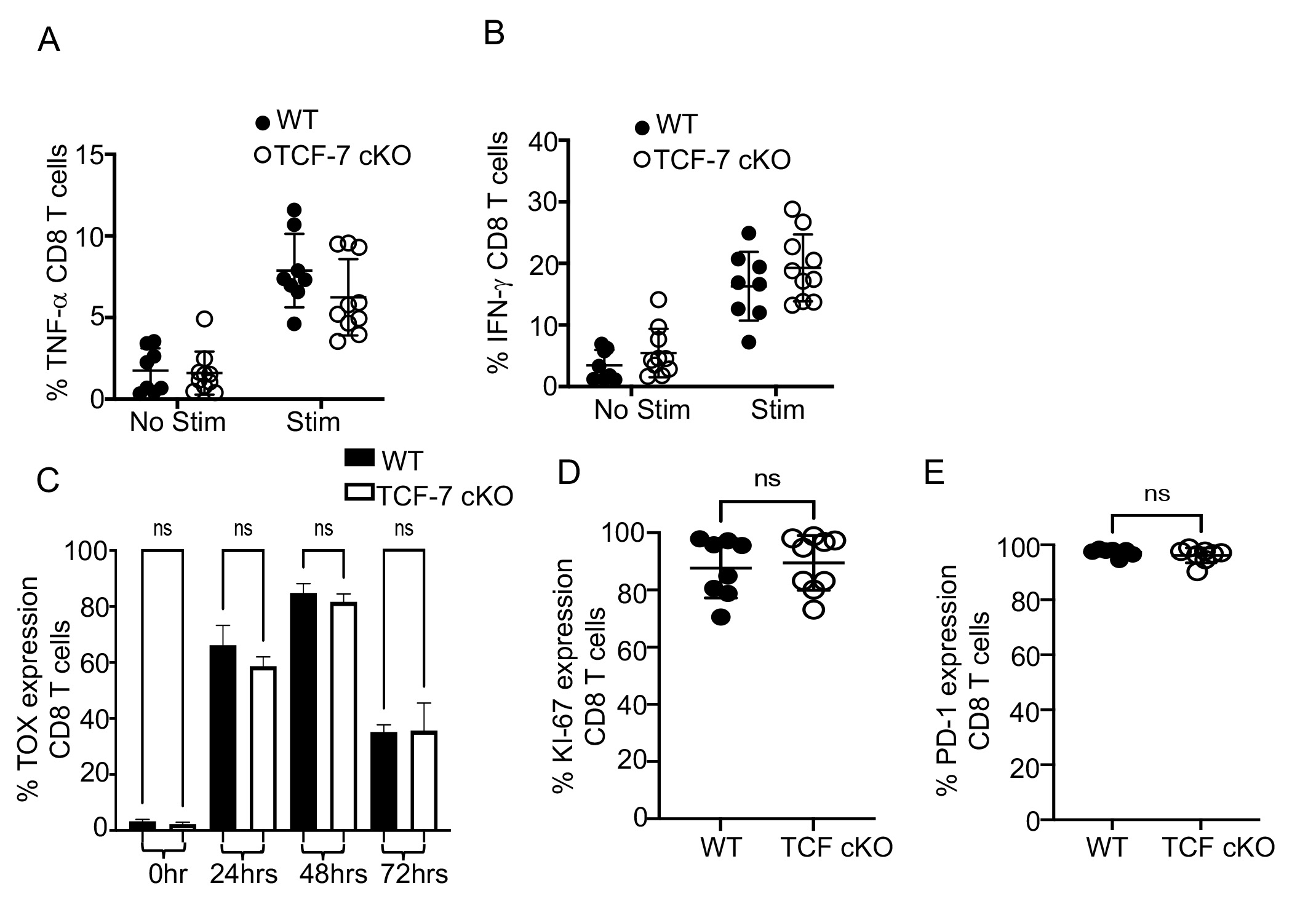
